# Cleavage of the catalytic intermediate fine-tunes cGAS signalling

**DOI:** 10.1101/2025.03.04.641389

**Authors:** Yan Yan, Xiang Li, Shuaihua Zhang, Xinyuan Zhao, Bingbing Yu, Xinwei Wang, Si Ai, Zhonghan Chen, Haoyu Wang, Shu Meng, Longfei Wang, Bin Zhu

**Affiliations:** Key Laboratory of Molecular Biophysics, the Ministry of Education, College of Life Science and Technology, Huazhong University of Science and Technology; Wuhan, 430074, China; Department of Cardiology, Taikang Center for Life and Medical Sciences, Zhongnan Hospital of Wuhan University, School of Pharmaceutical Sciences, Wuhan University; Wuhan, 430074, China; Department of Basic Science Research, Guangzhou National Laboratory, Guangzhou, Guangdong, 510005, China; Shenzhen Huazhong University of Science and Technology Research Institute; Shenzhen, 518063, China

## Abstract

Excessive activation of cyclic GMP–AMP synthase (cGAS) drives inflammatory and autoimmune diseases, but complete inhibition of cGAS signaling can compromise host defence. Here we show that cGAS signaling can be tuned through a previously unrecognized regulatory checkpoint. Bacterial cGAS generates cGAMP through a two-step mechanism involving release and rebinding of a 2′–5′-linked linear dinucleotide intermediate. We identify a CBASS-associated exonuclease (Exo) that selectively degrades this catalytic intermediate, establishing a threshold for cGAS activation while preserving immune function. This thresholding mechanism is conserved across domains of life: Exo suppresses human cGAS activity in vitro and in mammalian cells and attenuates cGAS-driven pathology in vivo. Our findings identify the catalytic intermediate surveillance as a conserved strategy for safeguarding cGAS signaling.

## Main Text

Cyclic GMP–AMP synthase (cGAS) is a central initiator of innate immune responses to infection-associated signals (*1–6*). Upon activation, cGAS catalyses the production of cyclic GMP–AMP, triggering signalling cascades that can culminate in irreversible cellular shutdown or programmed cell death (*7–17*). While this potent response is essential for host defence, inappropriate or excessive cGAS activation is deleterious and has been linked to autoinflammatory and autoimmune disease (*18–23*). These opposing demands impose a fundamental constraint on cGAS-family enzymes: signalling must be tightly restrained under basal conditions, yet capable of rapid and robust activation upon infection.

Across domains of life, cGAS-family signalling is antagonized through several conserved mechanisms. These include degradation of the cyclic messenger (*24–26*), sequestration of signalling molecules (*27*), and direct inhibition of the synthase (*28*). Although effective at silencing immune activation, such strategies function primarily as binary “off switches” and are poorly suited to modulating signalling output in a graded or conditional manner. How cGAS activity is intrinsically tuned to prevent spurious activation while preserving responsiveness remains unclear.

Here we define a regulatory mechanism that operates at the level of cGAS catalysis. We show that bacterial cGAS synthesizes cGAMP through a discontinuous two-step mechanism that generates a diffusible 2′–5′-linked intermediate. A family of CBASS-associated exonucleases (Exo) selectively degrades this intermediate, rather than the final cyclic product, thereby intercepting signalling before completion. This intermediate-targeting activity imposes an activation threshold that suppresses basal signalling while permitting robust cGAMP production once catalytic flux exceeds Exo capacity.

Structural and biochemical analyses reveal that Exo enzymes are specialized DEDDh family nucleases that selectively recognize 2′–5′ phosphodiester linkages, defining a distinct evolutionary lineage adapted for catalytic intermediate targeting. This regulatory strategy is conserved across domains of life: Exo also degrades the catalytic intermediate of human cGAS, suppresses cGAS–STING signalling in mammalian cells, and attenuates cGAS-driven pathology in vivo. Together, our findings establish catalytic intermediate targeting as a checkpoint mechanism in cyclic oligonucleotide signalling and reveal selective degradation of catalytic intermediates as a conserved strategy for regulating innate immunity.

### Exo targets a 2′–5′-linked catalytic intermediate during cGAS signalling

Survey of publicly available CBASS operons revealed that ∼4% encode Exo homologues, most frequently associated with type II and type III systems (*11, 29*) (Fig. 1A). Sequence analysis indicated that these proteins harbor conserved motifs characteristic of the DEDDh family of exonucleases (Fig. 1B). However, in contrast to canonical DEDDh nucleases such as human TREX1, *Serratia marcescens* Exo (*Sm*Exo) exhibited no detectable nuclease activity toward a wide range of DNA or RNA substrates, including single- and double-stranded nucleic acids and DNA–RNA hybrids, even in the presence of the CBASS signalling molecule 3′,2′-cGAMP (Fig. 1C and Fig. S1A). Electrophoretic mobility shift assays further showed no measurable substrate binding (Fig. S1B).

**Fig. 1.**
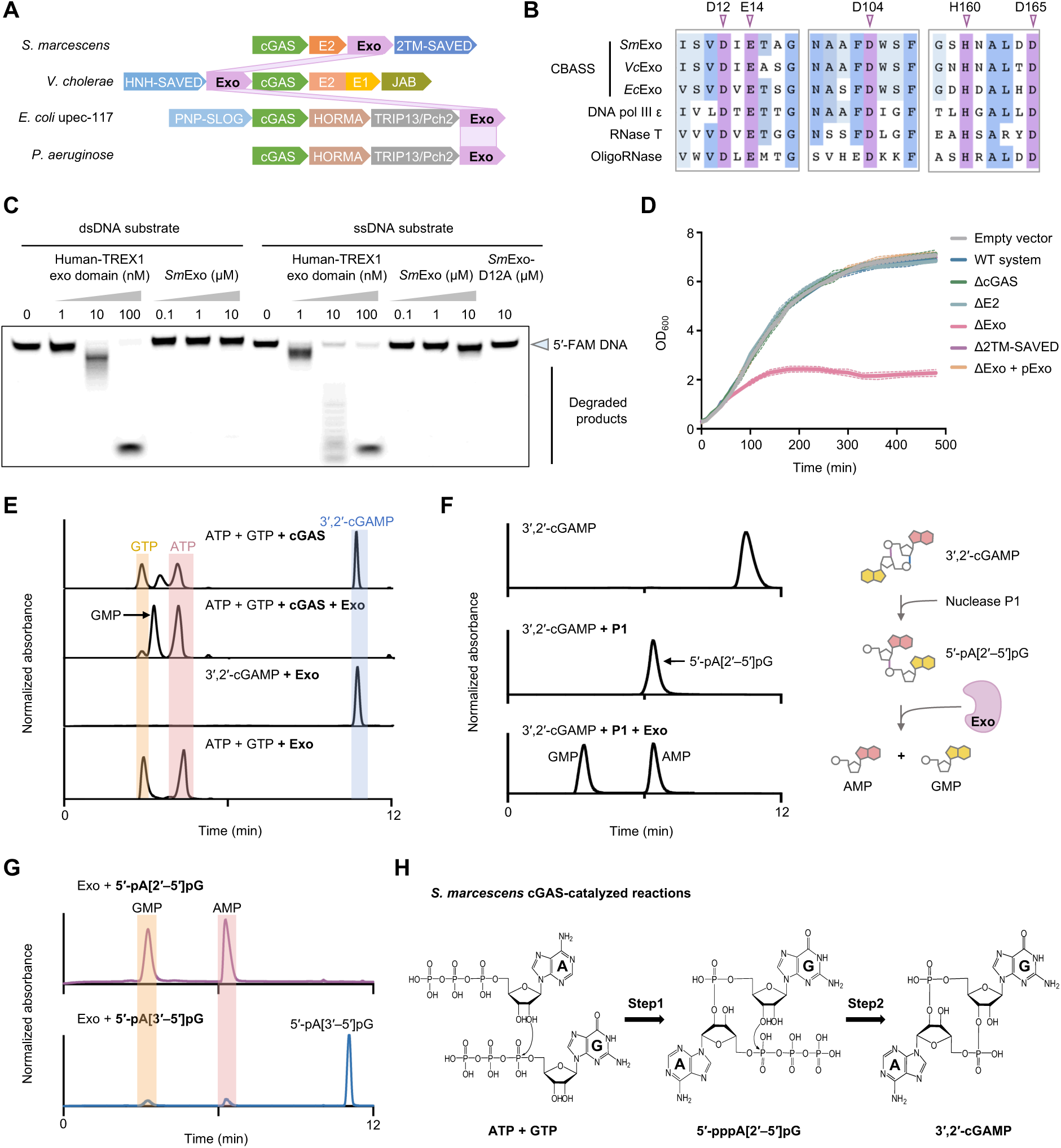
Exo selectively targets a 2′–5′-linked catalytic intermediate of cGAS. (**A**) Domain organization of representative CBASS operons from *Serratia marcescens*, *Vibrio cholerae*, *Escherichia coli* upec-117 and *Pseudomonas aeruginosa*, highlighting Exo homologues in a subset of type II and type III systems. Approximately 4% of annotated CBASS operons encode Exo. (**B**) Sequence alignment of CBASS-associated Exo proteins with canonical DEDDh family exonucleases, showing conservation of the catalytic DEDDh motif. (**C**) Nuclease activity assay using 5′-FAM-labelled dsDNA and ssDNA substrates incubated with *Sm*Exo or the human TREX1 exonuclease domain at the indicated concentrations. (**D**) Bacterial growth curves showing cytotoxicity upon deletion of *exo* and rescue by complementation. (**E**) HPLC analysis of in vitro reactions showing inhibition of 3′,2′-cGAMP synthesis by *Sm*Exo. *Sm*Exo does not hydrolyse 3′,2′-cGAMP or ATP and GTP alone but promotes conversion of GTP to GMP in the presence of active cGAS. (**F**) HPLC analysis showing hydrolysis of 5′-pA[2′–5′]pG, generated by nuclease P1 cleavage of 3′,2′-cGAMP, into AMP and GMP by Exo. (**G**) HPLC analysis comparing Exo activity toward 5′-pA[2′–5′]pG and 5′-pA[3′–5′]pG. (**H**) Schematic of the two-step pathway for 3′,2′-cGAMP synthesis via a 5′-pppA[2′–5′]pG intermediate.

Despite this apparent lack of canonical nuclease activity, deletion of *exo* in *S. marcescens* resulted in severe cytotoxicity, consistent with hyperactivation of the CBASS pathway (Fig. 1D). This phenotype depended on cGAMP signalling and activation of the membrane-disrupting effector 2TM-SAVED (*15–17*), indicating that Exo functions as a negative regulator of cGAS. Consistently, addition of Exo to in vitro reactions strongly inhibited synthesis of 3′,2′-cGAMP by cGAS (Fig. 1E). Importantly, Exo prevented further accumulation of cGAMP when added to ongoing reactions without reducing preformed product (Fig. S1C), suggesting that Exo acts upstream of final product formation.

Exo did not hydrolyze 3′,2′-cGAMP, nor did it exhibit nucleoside triphosphatase activity toward ATP or GTP alone (Fig. 1E), ruling out direct degradation of substrates or products. However, in the presence of active cGAS, Exo induced conversion of GTP to GMP (Fig. 1E). This observation cannot be explained by direct substrate or product turnover, suggesting that Exo acts on a reaction intermediate generated during catalysis.

To further investigate this activity, we generated a defined dinucleotide substrate by selectively cleaving the 3′–5′ phosphodiester bond of 3′,2′-cGAMP using nuclease P1, yielding 5′-pA[2′–5′]pG (Fig. 1F). Notably, Exo efficiently hydrolyzed this molecule into AMP and GMP in a divalent metal ion–dependent manner, with Mg²⁺ or Mn²⁺ supporting catalysis (Fig. 1F and Fig. S1D). In contrast, Exo displayed minimal activity toward the corresponding 3′–5′-linked analogue (Fig. 1G and Fig. S1E). These results reveal a previously unrecognized enzymatic activity within the DEDDh family: selective cleavage of 2′–5′ phosphodiester bonds.

This marked linkage selectivity raised the possibility that Exo targets a physiologically relevant 2′–5′-linked intermediate. Notably, synthesis of 3′,2′-cGAMP by cGAS has been reported to proceed through a sequential two-step mechanism involving formation of a linear 2′–5′-linked intermediate (*30*) (Fig. 1H). If Exo acts at the level of catalysis, such an intermediate must be generated and become accessible during the reaction. We therefore sought to determine whether this intermediate is produced and whether it is susceptible to Exo-mediated degradation.

Using non-hydrolysable nucleotide analogues to trap reaction intermediates, we found that replacing ATP with Ap(c)pp still permitted accumulation of a linear intermediate, whereas substitution of GTP with Gp(c)pp abolished product formation (Fig. 2A). These results demonstrate that *S. marcescens* cGAS generates a 5′-pppA[2′–5′]pG intermediate during cGAMP synthesis. Exo efficiently converted the 5′-pp(c)pA[2′–5′]pG intermediate into GMP and Ap(c)pp, indicating that it directly targets the catalytic intermediate rather than the final cyclic product.

**Fig. 2.**
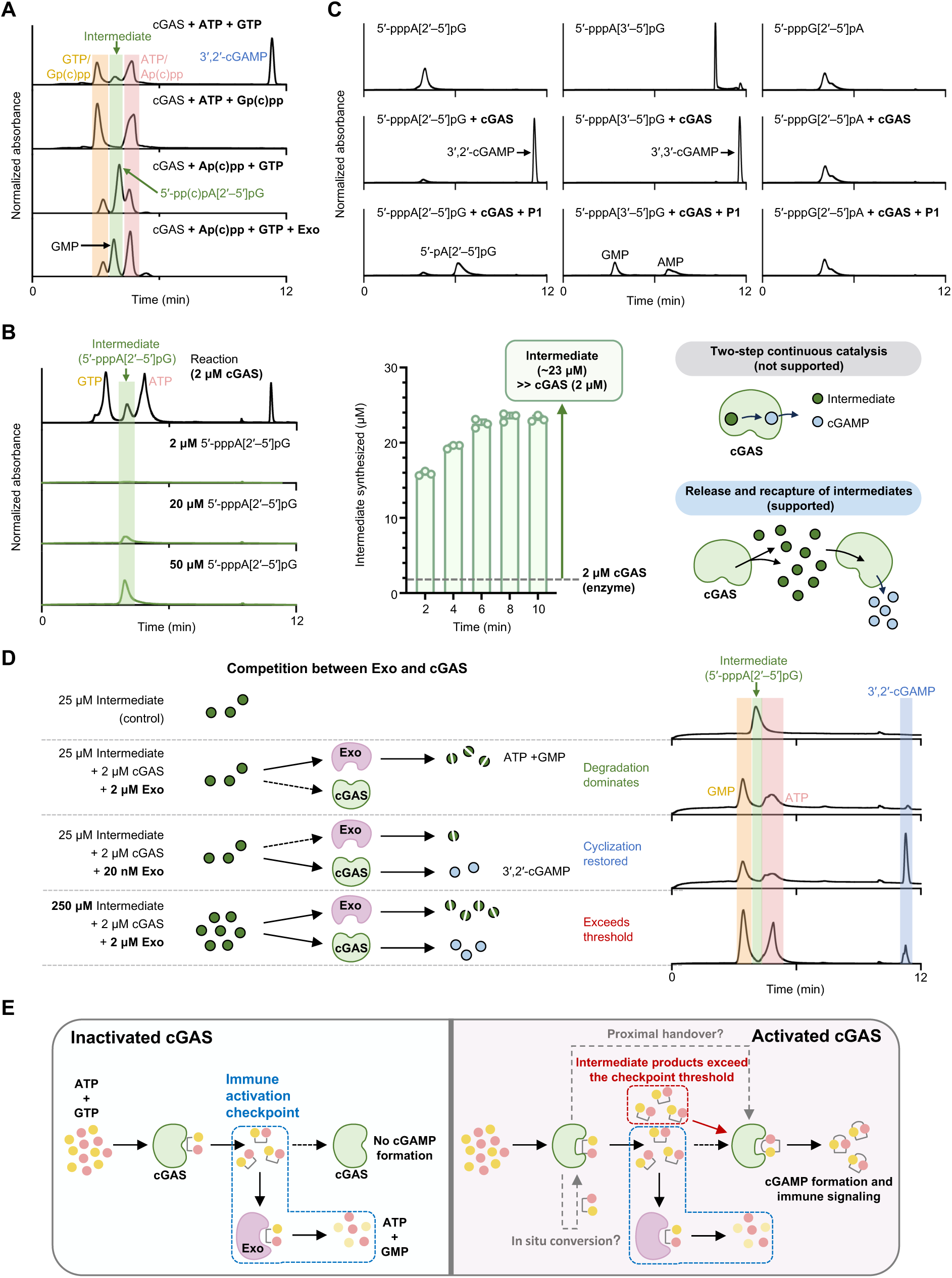
A diffusible catalytic intermediate enables threshold checkpoint control. (**A**) HPLC analysis of *S. marcescens* cGAS reactions in which ATP or GTP was replaced with non-hydrolysable analogues Ap(c)pp or Gp(c)pp. Addition of Exo results in degradation of the intermediate and production of GMP. (**B**) Left, HPLC detection of the cGAS-generated intermediate (5′-pppA[2′–5′]pG), identified by comparison with a synthetic standard and quantified using independently run standards. Middle, quantification of intermediate accumulation over time. Data are mean ± s.d., n = 3 independent experiments. Right, model illustrating release of the intermediate into solution rather than retention in a processive catalytic cycle. (**C**) HPLC analysis demonstrating that *S. marcescens* cGAS cyclizes 5′-pppA[2′–5′]pG and 5′-pppA[3′–5′]pG through formation of a 3′–5′ phosphodiester bond, whereas 5′-pppG[2′–5′]pA is not a substrate for cyclization. (**D**) Left, schematic of reactions containing cGAS, Exo and the linear intermediate under the indicated conditions. Right, corresponding HPLC analysis showing depletion of intermediate and accumulation of cyclization products. (**E**) Model of checkpoint control. Under low cGAS activity, the intermediate is degraded by Exo following release, preventing re-engagement with cGAS. Increased cGAS activity allows intermediate accumulation above a threshold, enabling productive cyclization and signalling.

### A discontinuous catalytic mechanism exposes a checkpoint for Exo-mediated regulation

We next sought to determine how Exo accesses the catalytic intermediate generated by cGAS. One possibility was that Exo directly interacts with cGAS to intercept the intermediate during catalysis. However, pull-down assays failed to detect any interaction between Exo and cGAS, even under active reaction conditions (Fig. S2, A to C), arguing against a substrate-channeling mechanism.

We therefore considered an alternative model in which the intermediate is released into solution, thereby becoming accessible to Exo. Quantitative analysis revealed that the concentration of the 5′-pppA[2′–5′]pG intermediate accumulated to ∼23 μM in reactions containing only 2 μM cGAS. This concentration substantially exceeds that of the enzyme (Fig. 2B). These observations indicate that the intermediate is released into solution rather than retained within a single catalytic cycle. However, these excess intermediates were eventually all turned to cyclic products (Fig. S3A). Together, these findings support a discontinuous catalytic mechanism in which the intermediate is transiently released and must subsequently rebind for cyclization (Fig. 2B).

Such a mechanism renders the intermediate accessible to competing enzymes, including Exo. Consistent with this interpretation, the catalytic intermediate was not intrinsically committed to a single reaction outcome. Although *Serratia marcescens* cGAS predominantly synthesizes 3′,2′-cGAMP via a 5′-pppA[2′–5′]pG intermediate, it was also capable of utilizing both 5′-pppA[2′–5′]pG and 5′-pppA[3′–5′]pG intermediates to form cyclic products through a second 3′–5′ phosphodiester bond, yielding 3′,2′-cGAMP and 3′,3′-cGAMP, respectively (Fig. 2C). However, this cyclization reaction remained dependent on nucleotide identity, as the reciprocal intermediate 5′-pppG[2′–5′]pA failed to undergo productive cyclization. These findings indicate that the intermediate is released as a chemically flexible species that can be re-engaged by cGAS, further supporting the existence of an accessible intermediate pool.

To test whether Exo competes with cGAS for this intermediate pool, we reconstituted reactions containing defined amounts of cGAS, intermediate and Exo. At low intermediate concentrations and equimolar cGAS and Exo, the intermediate was preferentially degraded, preventing cGAMP production. Increasing intermediate concentration or reducing Exo levels shifted the reaction toward productive cGAMP synthesis (Fig. 2D). Similar behavior was observed upon varying Exo concentrations, with high Exo levels favoring suppression and lower Exo levels restoring cGAMP production (Fig. S3B). Consistent with this competitive model, titration of cGAS relative to Exo revealed a sharp transition from complete suppression to robust cGAMP synthesis as cGAS activity increased (Fig. S3C). These results indicate that Exo and cGAS compete for a shared pool of catalytic intermediate. This competition produces a sharp transition in signalling output, consistent with a threshold mechanism in which Exo establishes a catalytic checkpoint that must be overcome for signalling to proceed.

Together, these findings establish catalytic intermediate targeting as a regulatory principle in cyclic oligonucleotide signalling. Exo imposes a catalytic checkpoint by selectively degrading a diffusible 2′–5′-linked intermediate. Under basal conditions, Exo limits intermediate availability to prevent spurious signalling, whereas increased cGAS activity overcomes this constraint to enable productive cGAMP synthesis (Fig. 2E).

To assess the physiological relevance of this catalytic intermediate checkpoint, we examined the consequences of Exo activity in vivo. Co-expression of cGAS and the membrane-disrupting effector 2TM-SAVED resulted in severe cytotoxicity, whereas expression of either component alone had no effect. Co-expression of Exo suppressed this phenotype (Fig. S4A). Consistently, disruption of Exo activity impaired CBASS-mediated resistance to phage infection, phenotypes that were rescued by reintroduction of Exo, as shown by phage T4 infection assays (Fig. S4B). These findings demonstrate that Exo-mediated control of the catalytic intermediate is required to maintain a threshold checkpoint for CBASS activation in vivo.

### A structurally encoded and evolutionarily conserved mechanism for 2′–5′ linkage selectivity

Phylogenetic analysis revealed that CBASS-associated Exo homologues form a distinct clade within the DEDDh exonuclease family, including representatives from *Serratia marcescens* (*17*), *Vibrio cholerae* (*29*) and *Escherichia coli* upec-117 (*31*), clearly segregating from canonical 3′–5′ exonucleases with established nucleic acid degradation activity (Fig. 3A and Fig. S5). This separation suggests that CBASS-associated Exo proteins have undergone functional specialization.

**Fig. 3.**
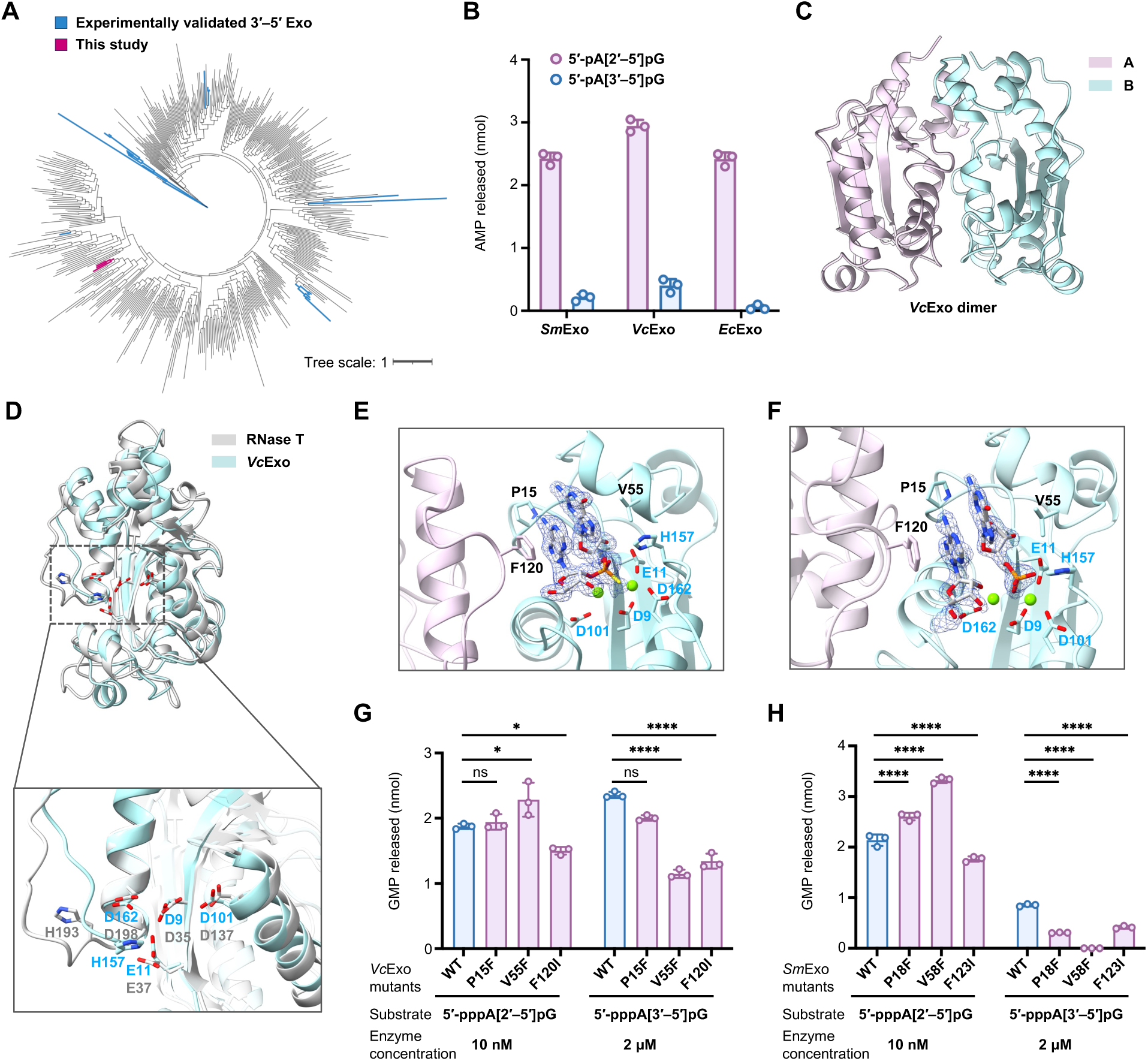
Structural basis of selective 2′–5′ linkage targeting by Exo. (**A**) Phylogenetic tree of DEDDh family exonucleases showing that CBASS-associated Exo proteins form a distinct clade. Exo homologues analysed in this study are highlighted in red, and experimentally validated 3′–5′ exonucleases are shown in blue. (**B**) Quantification of hydrolysis of 5′-pA[2′–5′]pG and 5′-pA[3′–5′]pG by Exo homologues from *S. marcescens*, *V. cholerae* and *E. coli* upec-117. Data are mean ± s.d., n = 3 independent experiments. (**C**) Crystal structure of *Vibrio cholerae* Exo (*Vc*Exo) shown as a homodimer. (**D**) Structural comparison of *Vc*Exo with the canonical DEDDh exonuclease RNase T from *P. aeruginosa* (PDB: 2F96). (**E**) Structure of *Vc*Exo bound to a non-hydrolysable A[2′–5′]pG analogue A[2′–5′]psG, showing the substrate-binding pocket. Active-site residues and Mg²⁺ ions are indicated. (**F**) Structure of *Vc*Exo bound to the hydrolysable substrate A[2′–5′]pG. The substrate adopts a conformation similar to that of the non-hydrolysable analogue. (**G**) HPLC quantification of cleavage of 5′-pppA[2′–5′]pG and 5′-pppA[3′–5′]pG by wild-type *Vc*Exo and its mutants. Data are mean ± s.d., n = 3 independent experiments. (**H**) HPLC quantification of cleavage of 5′-pppA[2′–5′]pG and 5′-pppA[3′–5′]pG by wild-type *Sm*Exo and its mutants. Data are mean ± s.d., n = 3 independent experiments.

Across diverse bacterial species, CBASS-associated Exo homologues consistently displayed robust hydrolytic activity toward 2′–5′-linked dinucleotide substrates, while exhibiting minimal activity toward the corresponding 3′–5′-linked analogues (Fig. 3B and Fig. S6A).

Consistent with this conserved specificity, Exo homologues efficiently suppressed cGAMP synthesis and selectively degraded 2′–5′-linked intermediates, while showing limited activity toward 3′–5′-linked substrates and related signalling systems (Fig. S6, B to C). These findings establish selective 2′–5′ phosphodiesterase activity as a defining functional feature of this protein family.

Notably, the *Vibrio cholerae* CBASS system analysed here encodes a 3′,2′-cGAMP–producing cGAS (Fig. S6D), in contrast to canonical DncV-type systems that synthesize 3′,3′-cGAMP(*9, 32*), further underscoring the diversity of cyclic dinucleotide signalling pathways targeted by Exo.

To elucidate the structural basis for this unusual linkage selectivity, we determined the crystal structure of *Vibrio cholerae* Exo (*Vc*Exo). The structure revealed a *C*2-symmetric homodimeric architecture characteristic of DEDDh family nucleases (Fig. 3C). Structural comparison with the predicted structure of *Serratia marcescens* Exo (*Sm*Exo) and the crystal structure of *E. coli* upec-117 Exo (*31*) (*Ec*Exo) demonstrated a highly conserved overall fold across species (Fig. S7, A to C), indicating that CBASS-associated Exo proteins share a common structural framework. Comparison with *Pseudomonas aeruginosa* RNase T (PDB: 2F96) revealed a broadly similar catalytic core, albeit with distinct positioning of the conserved histidine residue within the DEDDh motif, suggesting mechanistic divergence (Fig. 3D).

To define how Exo recognizes 2′–5′-linked substrates, we solved crystal structures of *Vc*Exo in complex with both A[2′–5′]pG and its nonhydrolyzable phosphorothioate analogue at resolutions of 1.99Å and 1.96Å, respectively (Fig. 3, E and F). Both crystals belong to the *C*2 space group, with six molecules per asymmetric unit. However, due to crystal packing, only a subset (two for A[2′–5′]pG, three for A[2′–5′]psG) of these molecules have well-defined substrate density. Notably, A[2′–5′]pG is cleaved into adenosine (A) and guanosine 5′-phosphate (pG) in the crystal structure with *Vc*Exo, suggesting that A[2′–5′]pG is a substrate for Exo.

In both structures, the A[2′–5′]psG substrate and products adopt nearly identical conformations within the active site. Substrate binding is stabilized by π–π stacking interactions between the adenine and guanine bases, and the aromatic side chain of F120 from the opposing protomer. This triple-stacking constrains residues A and G, thereby positioning their 2′–5′ linkage for catalysis. This configuration is compatible with 2′–5′ linkages but would be sterically disfavoured for 3′–5′ linkages, providing a structural basis for the observed selectivity.

Additional interactions further position the substrate within the catalytic pocket. Two Mg²⁺ ions are coordinated by conserved residues (D9, E11, D101 and D162) together with a phosphate oxygen, consistent with a two-metal-ion catalytic mechanism. The guanine base forms hydrogen bonds with D49 and E51, while the ribose hydroxyl groups are positioned for interactions with E11 and N97. The conserved H157 residue is ideally positioned to activate a water molecule for nucleophilic attack on the phosphodiester bond, supporting its role as a catalytic switch. Consistent with this structural model, mutations within the conserved DEDDh motif abolished hydrolysis of 5′-pA[2′–5′]pG and eliminated Exo-mediated suppression of cGAMP synthesis in vitro (Fig. S8, A and B), confirming the essential role of the catalytic core.

Closer inspection of the substrate-binding groove revealed structural features that could bias Exo toward 2′–5′ linkages. To test this, we introduced targeted mutations designed to alter the geometry of the binding pocket. In *Vc*Exo, mutation V55F enhanced cleavage of the native 2′–5′-linked intermediate while reducing activity toward the 3′–5′-linked analogue, indicating a shift in linkage preference. Mutation P15F produced a similar but less pronounced effect. In contrast, mutation F120I reduced cleavage of both substrates, indicating a general catalytic defect rather than altered specificity (Fig. 3G and Fig. S8C).

To assess whether this mechanism is conserved, we introduced corresponding mutations into *Sm*Exo guided by sequence alignment and the predicted structural model. Mutations *Sm*Exo V58F and *Sm*Exo P18F similarly enhanced cleavage of 2′–5′-linked substrates while reducing activity toward 3′–5′ linkages, whereas mutation *Sm*Exo F123I impaired cleavage of both substrates (Fig. 3H and Fig. S8D). Additionally, mutation in the conserved DEDDh motif of *Sm*Exo abolished catalytic activity and eliminated its ability to suppress cGAMP synthesis (Fig. S8, E and F), further demonstrating conservation of the catalytic mechanism.

To further assess whether substrate recognition is determined by phosphodiester linkage rather than substrate class, we examined Exo activity toward 2′–5′-linked oligoadenylates, which function as signalling molecules in eukaryotic innate immunity (*6*). Exo efficiently cleaved a 2′–5′-linked oligoadenylate (2′–5′-AAAAA), as shown by HPLC analysis (Fig. S9A). Polyacrylamide gel electrophoresis further confirmed 3′-to-5′ exonucleolytic degradation of the 2′–5′-linked oligoadenylate substrate (Fig. S9B). These results indicate that Exo recognizes substrates primarily through 2′–5′ phosphodiester linkage geometry.

Together, these findings identify a structurally encoded and evolutionarily conserved substrate-recognition mechanism that enforces selective cleavage of 2′–5′ phosphodiester bonds. These results establish CBASS-associated Exo proteins as a distinct functional lineage within the DEDDh family consisting mostly of exonucleases that cleave 3′–5′ phosphodiester bonds in DNA or RNA, specialized for targeting 2′–5′-linked intermediates in cyclic oligonucleotide signalling.

### Exo targets human cGAS intermediates to impose a conserved catalytic checkpoint

Having established that Exo suppresses bacterial cGAS signalling by selectively degrading a 2′–5′-linked catalytic intermediate, we next asked whether this regulatory principle extends to eukaryotic cyclic GMP–AMP synthase. Human cGAS produces 2′,3′-cGAMP through a sequential two-step reaction that proceeds via a 5′-pppG[2′–5′]pA intermediate (Fig. 4A).

**Fig. 4.**
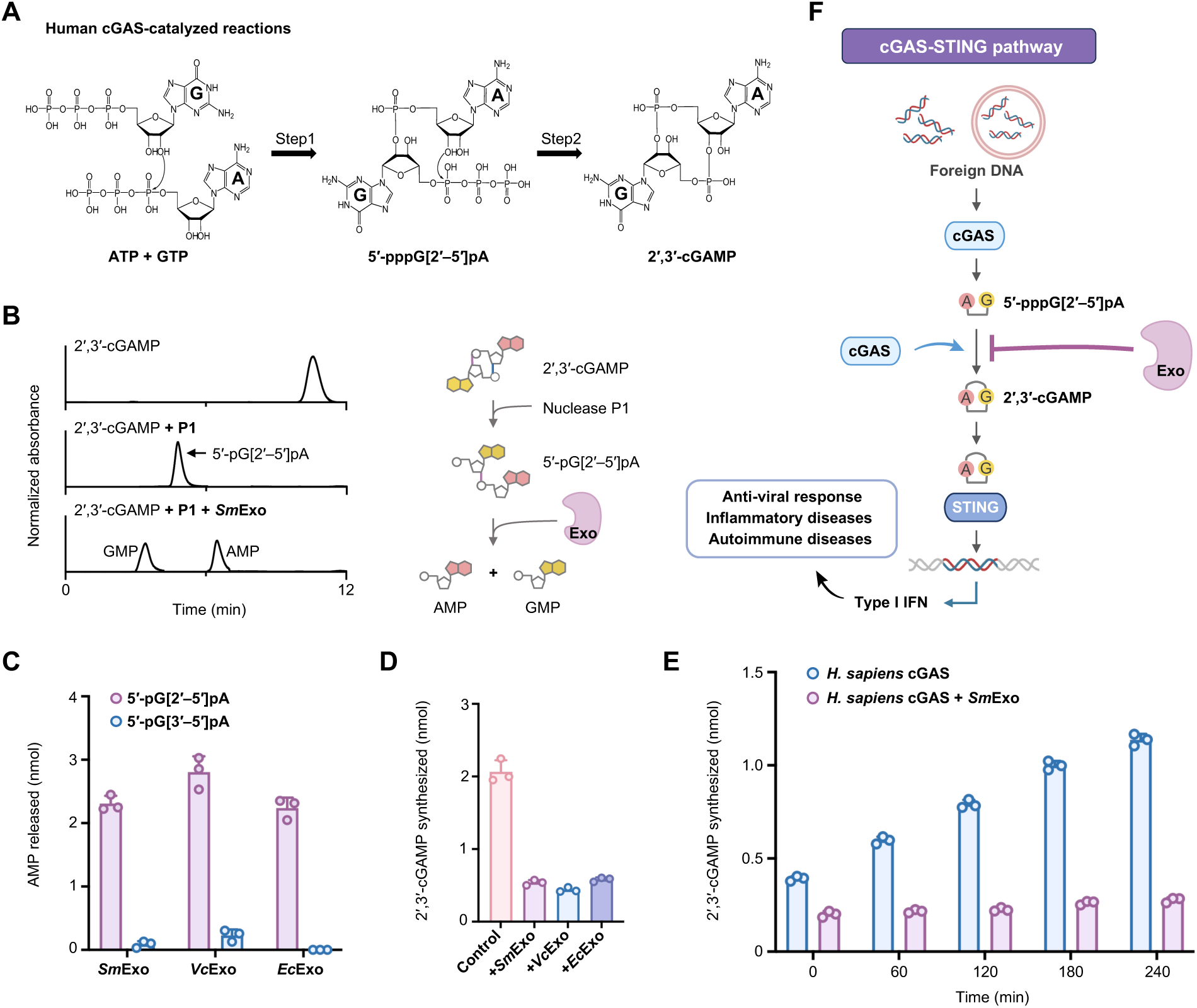
Exo targets the catalytic intermediate of human cGAS. (**A**) Schematic of human cGAS catalysis, showing formation of 2′,3′-cGAMP via a 5′-pppG[2′–5′]pA intermediate. (**B**) HPLC analysis showing hydrolysis of 5′-pG[2′–5′]pA, generated from 2′,3′-cGAMP, into AMP and GMP by *Sm*Exo. (**C)** Quantification of hydrolysis of 5′-pG[2′–5′]pA and 5′-pG[3′–5′]pA by Exo homologues from *S. marcescens*, *V. cholerae* and *E. coli* upec-117. Data are mean ± s.d., n = 3 independent experiments. (**D**) Quantification of 2′,3′-cGAMP synthesis by human cGAS in the presence or absence of Exo homologues. Data are mean ± s.d., n = 3 independent experiments. (**E**) Time-course analysis of 2′,3′-cGAMP accumulation in human cGAS reactions with or without Exo. Data are mean ± s.d., n = 3 independent experiments. (**F**) Model illustrating suppression of cGAS–STING signalling by Exo through degradation of the catalytic intermediate.

The shared use of a 2′–5′-linked catalytic intermediate raised the possibility that Exo might similarly target human cGAS at the level of catalysis.

To test this, we generated a defined dinucleotide substrate by selectively cleaving the 3′–5′ phosphodiester bond of 2′,3′-cGAMP using nuclease P1, yielding 5′-pG[2′–5′]pA (Fig. 4B). Exo efficiently hydrolyzed this molecule into AMP and GMP, demonstrating that it can directly target the catalytic intermediate generated during human cGAS activity. Notably,

Exo cleaved 5′-pG[2′–5′]pA with efficiency comparable to that observed for the bacterial analogue 5′-pA[2′–5′]pG, indicating that substrate recognition is largely independent of base identity and instead dictated by phosphodiester linkage geometry (Fig. S10A).

Consistent with this linkage-driven specificity, Exo homologues from multiple bacterial species robustly hydrolyzed 5′-pG[2′–5′]pA while exhibiting minimal activity toward the corresponding 3′–5′-linked substrate (Fig. 4C), demonstrating that recognition of the human cGAS intermediate is a conserved property of CBASS-associated Exo proteins.

We next examined the functional consequences of this activity in the context of human cGAS catalysis. Addition of Exo to in vitro reactions markedly suppressed accumulation of 2′,3′-cGAMP, and time-course analyses revealed that Exo limits signal accumulation over time (Fig. 4, D and E, and Fig. S10B).

Together, these findings demonstrate that Exo restrains cGAS–STING signalling by selectively degrading the catalytic intermediate of human cGAS, thereby imposing a conserved catalytic checkpoint on cyclic GMP–AMP production (Fig. 4F). This result extends the intermediate-targeting regulatory mechanism defined in bacterial CBASS systems to eukaryotic innate immunity and identifies catalytic intermediates as conserved control points in cyclic oligonucleotide signalling.

### Exo suppresses cGAS–STING signalling in mammalian systems and ameliorates inflammatory pathology

To determine whether the intermediate-targeting mechanism defined above operates in physiological settings, we examined the activity of Exo in mammalian cells and in vivo. We generated FLAG-tagged Exo mRNAs from multiple bacterial species and assessed their expression in human and murine cell lines. Detectable expression was observed only for *Vc*Exo, and was markedly enhanced by human codon optimization (Fig. 5A and Fig. S11, A and B). Codon-optimized *Vc*Exo was robustly expressed in RAW264.7 macrophages for more than 48 h without detectable cytotoxicity, even at high doses (Fig. 5, B and C, Fig. S11C), and was therefore used for subsequent experiments.

**Fig. 5.**
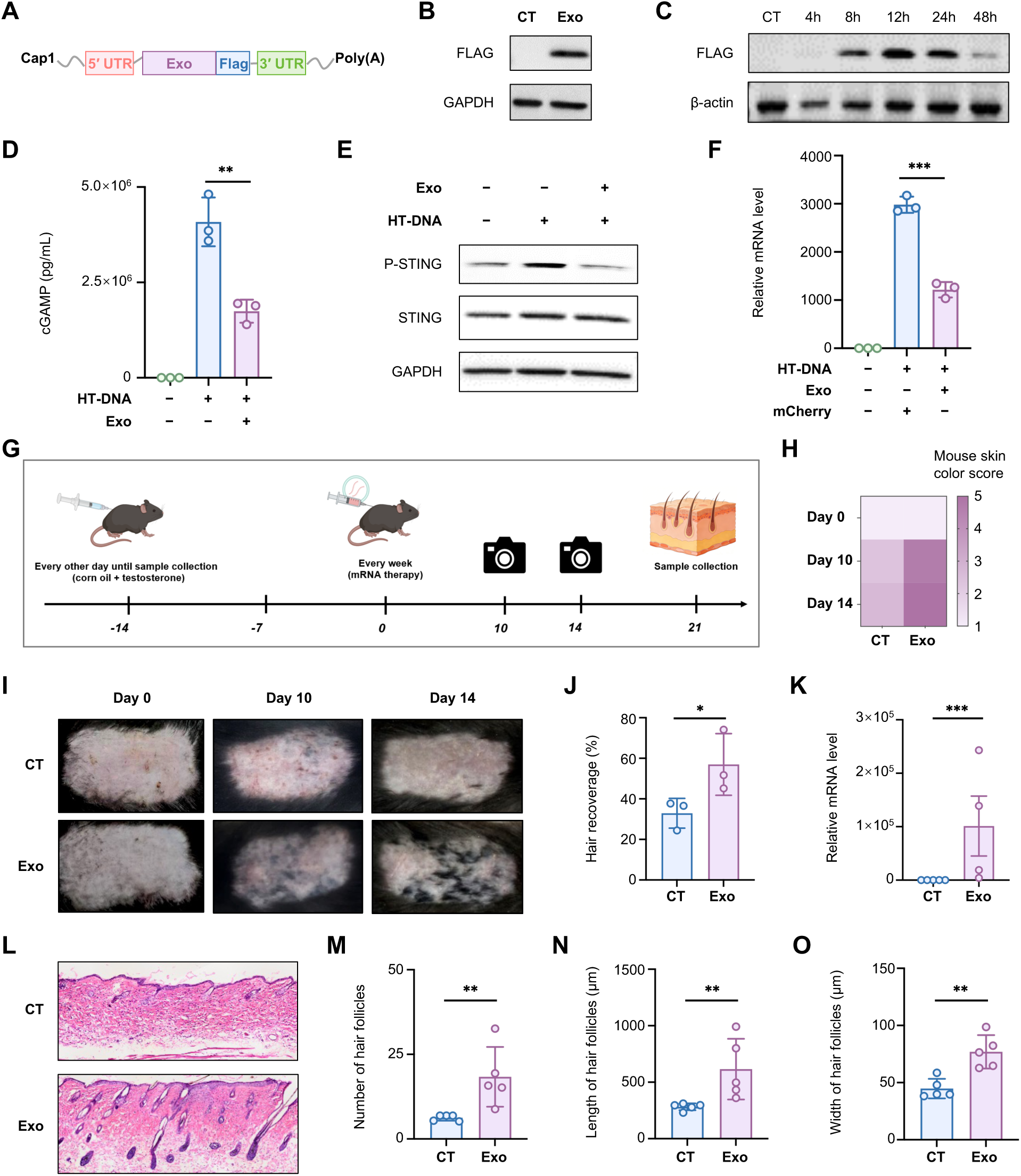
Exo imposes a threshold checkpoint on cGAS–STING signalling in mammalian systems. (**A**) Schematic of the codon-optimized, FLAG-tagged Exo mRNA construct used for mammalian expression. (**B**) Immunoblot analysis of Exo expression in RAW264.7 macrophages following mRNA transfection. (**C**) Time-course of Exo expression over 48 h. (**D**) Intracellular cGAMP levels in RAW264.7 cells pretreated with Exo mRNA and stimulated with HT-DNA. Data are mean ± s.d., n = 3 independent experiments. (**E**) Immunoblot analysis of STING phosphorylation under the indicated conditions. (**F**) Quantification of *Ifnb* induction. Data are mean ± s.d., n = 3 independent experiments. (**G**) Schematic of LNP-mediated delivery of Exo mRNA in a testosterone-induced mouse model of androgenetic alopecia. (**H-J**) Representative images and quantification of hair regrowth following treatment with Exo mRNA or control. (**K**) Quantification of Exo mRNA persistence in dorsal skin. Data are mean ± s.d., n = 5 mice per group. (**L-O**) Histological analysis of dorsal skin sections showing hair follicle number and size. Data are mean ± s.d., n = 5 mice per group.

Pretreatment of RAW264.7 cells with Exo mRNA substantially reduced intracellular cGAMP accumulation following HT-DNA transfection (Fig. 5D). Consistent with reduced availability of the cyclic dinucleotide signal, Exo expression suppressed STING phosphorylation and induction of interferon-β (Fig. 5, E and F). Importantly, mutations of conserved catalytic residues largely abolished these inhibitory effects (Fig. S11D), demonstrating that suppression of cGAS–STING signalling depends on Exo enzymatic activity and is consistent with catalytic intermediate targeting.

We next assessed the in vivo consequences of Exo expression in a mouse model of androgenetic alopecia, in which chronic activation of the cGAS–STING pathway driven by mitochondrial DNA release impairs hair follicle regeneration. Mice treated with lipid nanoparticles encapsulating Exo mRNA exhibited an accelerated transition from telogen to anagen, as evidenced by progressive skin pigmentation and robust hair regrowth compared with control-treated animals (Fig. 5, G to J). Consistent with sustained local delivery, Exo mRNA remained detectable in dorsal skin several days after administration (Fig. 5K).

Histological analysis confirmed enhanced follicular regeneration, with increased follicle number and size in Exo-treated animals (Fig. 5, L to O). No histopathological abnormalities were observed in major organs (Fig. S11E), indicating a favourable safety profile under these conditions.

Together, these findings demonstrate that Exo-mediated targeting of the cGAS catalytic intermediate is sufficient to suppress cGAS–STING signalling in mammalian cells and to ameliorate cGAS-driven pathology in vivo. These results establish catalytic intermediate targeting as a mechanistically grounded strategy for modulating innate immune signalling and highlight its potential for therapeutic intervention.

## Discussion

Our study identifies Exo as a critical regulator of cyclic oligonucleotide signalling in CBASS and uncovers a previously unrecognized mode of cGAS control. By selectively degrading a transient catalytic intermediate, Exo imposes an activation threshold that restrains signalling under basal conditions while permitting rapid amplification once enzymatic flux exceeds its capacity. This mechanism ensures that abortive infection is engaged only when the level of threat surpasses a defined threshold, thereby balancing antiviral defence with the substantial fitness costs of self-destruction.

Mechanistically, Exo operates through a mode of regulation distinct from previously described strategies. We detect no stable interaction between Exo and cGAS, arguing against direct enzyme sequestration or substrate channeling. Instead, our data support a model in which a linear catalytic intermediate becomes transiently accessible between successive phosphodiester bond–forming steps, allowing Exo to intercept the reaction prior to cyclization (Fig. 2B).

This framework resolves a long-standing paradox in cGAS enzymology. Previous studies of eukaryotic cGAS have proposed that linear intermediates dissociate from the active site prior to cyclization, with only a minor fraction undergoing immediate conversion to cGAMP (*5, 15–17, 33–35*). However, such dissociated intermediates would be expected to compete poorly with abundant ATP and GTP, leading to their accumulation. To reconcile this, it has been suggested that productive turnover may depend on inefficient rebinding, spatial confinement, or degradation by unidentified nucleases (*5, 15–17, 33–35*). Our results provide a direct resolution. We observe transient release of the linear intermediate during catalysis, and quantitative analysis shows that its concentration substantially exceeds that of cGAS, indicating that intermediates are released into solution rather than retained within a single catalytic cycle (Fig. 2B). Despite this, intermediates do not accumulate but remain at a steady-state level and are progressively depleted as cyclic product continues to accumulate. These observations support a model in which intermediate turnover limits accumulation and contributes to productive cGAMP synthesis, identifying intermediate dynamics as an important determinant of signalling output.

The physiological importance of this regulatory strategy is underscored by the consequences of Exo loss. Deletion of *exo* or disruption of its catalytic activity results in unrestrained CBASS activation, leading to cytotoxicity and impaired resistance to phage infection (Fig. 1D and Fig. S4, A and B). These findings demonstrate that precise control of intermediate turnover is essential not only to prevent inappropriate self-activation but also to optimize immune defence, highlighting the importance of threshold-based regulation in bacterial innate immunity.

At the molecular level, Exo defines a previously unrecognized specialization within the DEDDh exonuclease family. Whereas most members cleave canonical 3′–5′ phosphodiester bonds, Exo selectively hydrolyzes substrates containing 2′–5′ linkages (Fig. 1). This activity distinguishes Exo from structurally unrelated eukaryotic 2′–5′ phosphodiesterases involved in oligoadenylate turnover (*36–41*), which share similar substrate specificity but belong to distinct enzyme classes. Structural and biochemical analyses reveal how a conserved nuclease scaffold can be repurposed to recognize alternative phosphodiester geometries, and phylogenetic analysis further supports the emergence of a distinct lineage adapted for catalytic intermediate targeting.

These findings highlight the modularity of CBASS signalling systems. Diverse CBASS architectures encode cyclases that produce chemically distinct cyclic nucleotides, and the presence of Exo varies across subtypes (*11, 29*). The conserved preference of Exo for 2′–5′ linkages suggests that regulatory modules may co-evolve with specific signalling chemistries, raising the possibility that analogous mechanisms exist for other classes of cyclic nucleotide intermediates.

Beyond bacteria, we show that this intermediate-targeting strategy extends to eukaryotic innate immunity. Exo suppresses human cGAS activity by degrading its catalytic intermediate, thereby limiting activation of the cGAS–STING pathway in mammalian cells and ameliorating cGAS-driven pathology in vivo (*18, 42*) (Fig. 4 and 5). These findings indicate that catalytic intermediate targeting represents a conserved mechanism for tuning signalling output across domains of life.

Current strategies to therapeutically modulate cGAS activity focus largely on direct inhibition, including competition with substrates (*43, 44*), disruption of DNA binding (*45*), or suppression of cyclic dinucleotide production (*46*). While effective, such approaches risk broadly suppressing essential immune functions. In contrast, the mechanism described here operates by imposing a threshold at the level of catalysis, restraining basal signalling while preserving responsiveness. This framework suggests that targeting catalytic intermediates may provide a more balanced approach for modulating innate immune pathways.

Together, our work establishes catalytic intermediate targeting as a fundamental regulatory principle in cyclic oligonucleotide signalling. By demonstrating that innate immune responses can be controlled through selective turnover of enzymatic intermediates, we reveal a previously unappreciated layer of regulation that enables precise tuning of powerful second-messenger systems across domains of life.

## Acknowledgments

We thank all lab members for helpful discussions. We thank the staff at the beamline BL10U2 at Shanghai Synchrotron Radiation Facility.

## Funding

National Natural Science Foundation of China grant 32150009 (B.Z.) and 32571399 (L.F.W.).

Feng Foundation of Biomedical Research (B.Z.)

Billionhome Venture Capital (B.Z.)

National Key R&D Program of China grant 2025YFA0922101 (L.F.W.)

Large-scale Instrument and Equipment Sharing Foundation of Wuhan University (L.F.W.)

The Fundamental Research Funds for the Central Universities grant 2042025kf0012 (L.F.W.)

The Major Project of Guangzhou National Laboratory grant GZNL2025C02027 (S.M.)

## Author contributions

Conceptualization: B.Z., Y.Y., L.F.W., S.M.

Funding acquisition: B.Z., L.F.W., S.M.

Methodology: Y.Y., X.L., S.H.Z., X.Y.Z., B.B.Y., S.M., L.F.W., B.Z

Investigation: Y.Y., X.L., S.H.Z., X.Y.Z., B.B.Y., S.M., L.F.W., B.Z., S.A., X.W.W., Z.H.C., X.W.W.

Visualization: Y.Y., X.L., S.H.Z., X.Y.Z., S.M., L.F.W., B.Z.

Writing – original draft: Y.Y., X.L., S.H.Z., S.M., L.F.W., B.Z.

Writing – review & editing: B.Z., L.F.W., S.M., Y.Y., X.Y.Z.

## Data and materials availability

Structural coordinates and structure factors have been deposited in the Protein Data Bank under accession codes 22HD and 22HE. All other data are available in the main text or the supplementary materials. All materials are available from the corresponding authors upon reasonable request.

## Supplementary Materials

## Materials and Methods

### Plasmid construction

Protein sequences of *Serratia marcescens* cGAS (WP_016928966.1), *Vibrio cholerae* cGAS (WP_001901330.1), human cGAS (NP_612450.2), *S. marcescens* Exo (WP_016928968.1), *Escherichia coli* upec-117 Exo (WP_001258365.1), *V. cholerae* Exo (WP_001005971.1) were codon-optimized for *E. coli* expression and synthesized by GenScript. Genes were cloned into a custom pET28a-derived vector encoding an N-terminal hexahistidine tag. Untagged constructs used for pull-down assays were generated in pQE82L.

The exonuclease domain (residues 1–242) of human TREX1 (NP_338599.1) was codon-optimized for *E. coli* expression, synthesized by GenScript, and inserted into a custom pET28a-derived vector carrying an N-terminal His-SUMO tag.

For phage infection assays, the four-gene CBASS operon from *S. marcescens* (genomic coordinates 26,427–29,783) was cloned into pQE82L. Site-directed mutants were generated by overlap-extension PCR using the wild-type plasmid as template. All plasmids were propagated in *E. coli* DH5α and verified by Sanger sequencing.

### Protein expression and purification

Expression plasmids were transformed into *E. coli* BL21(DE3). Overnight starter cultures were grown in LB medium supplemented with appropriate antibiotics at 37 °C with shaking at 220 rpm. Cultures were diluted into fresh LB medium and grown to an OD600 of 0.8–1.0 before cooling to room temperature.

Protein expression was induced with 0.25 mM IPTG, and cultures were incubated overnight at 16 °C.

Cells were harvested by centrifugation and resuspended in lysis buffer containing 20 mM Tris-HCl, pH 8.0, 300 mM NaCl and 20 mM imidazole. Cells were lysed by sonication and clarified by centrifugation at 21,000 × g for 1 h at 4 °C. Clarified lysates were filtered through 0.45 µm filters and applied to Ni–NTA resin (Qiagen) using gravity-flow columns (Bio-Rad). Resin was washed sequentially with wash buffer (20 mM Tris-HCl, pH 8.0, 300 mM NaCl) supplemented with 50, 80 or 100 mM imidazole. Proteins were eluted in elution buffer containing 20 mM Tris-HCl, pH 8.0, 300 mM NaCl and 200 mM imidazole.

Eluted proteins were concentrated using centrifugal concentrators with 10 kDa or 30 kDa molecular-weight cutoffs (MilliporeSigma) and dialysed for ∼24 h at 4 °C against storage buffer containing 50 mM Tris-HCl, pH 7.5, 100 mM NaCl, 0.1 mM EDTA, 0.1% Triton X-100 and 50% glycerol. Proteins were stored at −20 °C.

The exonuclease domain of human TREX1 (residues 1–242), used as a positive control in nuclease assays, was purified using a modified protocol. Protein expression was induced with 0.4 mM IPTG and cultures were incubated overnight at 16 °C. Cells were lysed in buffer containing 20 mM HEPES, pH 7.5, 400 mM NaCl, 10% glycerol, 30 mM imidazole and 1 mM TCEP. Following Ni–NTA affinity purification, the His–SUMO tag was removed by incubation with ULP1 protease (∼250 µg) during overnight dialysis against 20 mM HEPES, pH 7.5, 150 mM NaCl and 1 mM TCEP at 4 °C. Untagged TREX1 was further purified by heparin affinity chromatography (HiTrap Heparin HP, Cytiva) using a linear gradient of 150–1,000 mM NaCl, followed by size-exclusion chromatography on a Superdex 200 Increase 10/300 GL column (Cytiva) equilibrated in 20 mM HEPES, pH 7.5, 250 mM NaCl and 1 mM TCEP. Peak fractions were concentrated to ∼10 mg mL^−1^, flash-frozen in liquid nitrogen and stored at −80 °C.

Protein concentrations were determined using Bradford assays (Bio-Rad) with bovine serum albumin as standard. Protein purity was assessed by Coomassie blue-stained SDS–PAGE and routinely exceeded 90%.

### Pull-down assay

His-tagged cGAS and untagged Exo proteins were expressed separately in *E. coli* and subjected to Ni–NTA pull-down assays. Clarified lysates were prepared by sonication and centrifugation as described above. Unless otherwise indicated, wash steps were performed using buffer supplemented with 20 mM imidazole.

For co-loading experiments, clarified cGAS and Exo lysates were mixed at a 1:1 ratio (total volume, 20 mL) before application to a 3 mL Ni–NTA affinity column. Flow-through fractions, low-imidazole wash fractions and 200 mM imidazole elution fractions were collected for analysis.

For sequential-loading experiments, clarified cGAS lysate was first applied to the Ni–NTA column. After collection of the initial flow-through and wash fractions, clarified Exo lysate was applied to the same column. A second set of flow-through and wash fractions, together with the final elution fraction, was subsequently collected.

As expression and solubility controls, Exo samples collected before induction, after induction, and from soluble and insoluble fractions following cell lysis were analysed by SDS–PAGE.

To investigate cGAS–Exo complex formation, gel filtration analyses were performed using a Superdex™ 200 Increase column (Cytiva). Samples (500 µL) contained: Exo alone (20 µM), cGAS alone (10 µM), Exo plus cGAS (20 µM and 10 µM, respectively), or Exo plus cGAS supplemented with 50 µM ATP and 50 µM GTP. Samples were prepared in buffer containing 50 mM Tris-HCl, pH 7.5, 100 mM NaCl, 2.5 mM MnCl2 and 1 mM DTT. For nucleotide-containing samples, ATP and GTP were additionally included in the running buffer. Mixed samples were incubated overnight on ice before injection.

### Nuclease assays

Unless otherwise stated, nuclease assays were performed in assay buffer containing 50 mM Tris-HCl, pH 7.5, 100 mM NaCl, 2.5 mM MnCl_2_ and 1 mM DTT.

To assess exonuclease activity, 10 µL reactions containing 1 µM nucleic acid substrate were supplemented with recombinant enzymes at the indicated concentrations. Human TREX1 exonuclease domain was used at 0, 1, 10 or 100 nM as a positive control. *S. marcescens* Exo (*Sm*Exo) was tested at 0.1, 1 or 10 µM, and the catalytic mutant *Sm*Exo-D12A was included at 10 µM as an inactive control. Substrates synthesized by GenScript included ssDNA (5′-FAM-CAATACCATATGACAGCTTAAATGTTTCCT-3′) and dsDNA of identical sequence.

Reactions were incubated at 37 °C for 10 min and terminated by addition of an equal volume of 2× RNA loading dye (New England Biolabs), followed by heating at 85 °C for 2 min and immediate cooling on ice. Products were resolved by 20% denaturing PAGE containing 8 M urea.

To evaluate substrate specificity in the presence of 3′,2′-cGAMP, reactions contained 2 µM nucleic acid substrate, 2 µM Exo and, where indicated, 2 µM 3′,2′-cGAMP. Substrates included dsDNA (Cas9 target fragment), ssDNA (5′-GCATAAACCTCATAATTAAAAATTTATCTAAGTCACTAACTC-3′), and corresponding DNA–RNA hybrid, dsRNA and ssRNA substrates. Reactions were incubated at 37 °C for 30 min and terminated by addition of EDTA-containing loading dye (final EDTA concentration, 20 mM). Products were analysed by agarose or polyacrylamide gel electrophoresis as appropriate.

### Electrophoretic mobility shift assays

Nucleic acid substrates (as above) were incubated with 2 µM Exo and/or 2 µM 3′,2′-cGAMP in assay buffer at 37 °C for 30 min. Reactions were terminated by addition of EDTA-containing loading dye. Samples were analysed using native agarose gels or native PAGE.

### Bacterial growth assays

Wild-type or mutant E2-CBASS operons were transformed into *E. coli* BL21. Cultures were grown at 37 °C to an OD600 of ∼0.2 before induction with 5 µM IPTG. Cells were incubated at 37 °C with shaking at 200 rpm, and OD600 was measured at the indicated time points.

### CD-NTase activity assays

CD-NTase reactions (20 µL) were performed in assay buffer supplemented with 250 µM ATP and GTP (New England Biolabs). Unless otherwise indicated, reactions contained 2 µM bacterial cGAS and 2 µM Exo. Human cGAS reactions were performed using 5 µM enzyme supplemented with 1 µM salmon sperm DNA (Sigma-Aldrich). Reactions were incubated at 37 °C for 0.5–2 h, heat-inactivated at 80 °C for 10 min and analysed by HPLC.

### Phosphodiesterase activity assays

Exo phosphodiesterase activity assays were performed in 20 µL reactions containing assay buffer, 250 µM substrate and 2 µM Exo. Substrates included 5′-pA[2′–5′]pG generated by nuclease P1 digestion of 3′,2′-cGAMP, 5′-pG[2′–5′]pA generated from 2′,3′-cGAMP. Reactions were incubated at 37 °C for 30 min and terminated by heating at 80 °C for 10 min before HPLC analysis.

For nuclease P1 treatment, reaction products were incubated with 1 µL nuclease P1 (New England Biolabs) for 1 h at 37 °C.

To assess phosphodiester bond selectivity, reactions containing 250 µM substrate were supplemented with wild-type or mutant Exo proteins. For 5′-pppA[2′–5′]pG (Biolog), reactions contained 10 nM enzyme and were incubated for 15 min at 37 °C. For 5′-pppA[3′–5′]pG (Biolog), reactions contained 2 µM enzyme and were incubated for 30 min at 37 °C. Reactions were terminated by heating at 80 °C for 10 min before HPLC analysis.

Cleavage of 2′–5′ oligoadenylates was analysed by HPLC and fluorescence gel assays. HPLC assays were performed in 20 µL reactions containing assay buffer, 40 µM 2′–5′-linked pentadenylate (2′–5′-AAAAA) and 10 µM Exo. Reactions were incubated at 37 °C for 30 min, heat-inactivated at 80 °C for 10 min and analysed by HPLC. Products were quantified by peak integration.

Fluorescence assays were performed in 10 µL reactions containing assay buffer, 5 µM 5′-FAM- or 3′-FAM-labelled 2′–5′-AAAAA substrate and 10 µM Exo. Reactions were incubated at 37 °C for 30 min and terminated by addition of an equal volume of 2× RNA loading dye (New England Biolabs), followed by heating at 85 °C for 2 min and immediate cooling on ice. Products were resolved by denaturing PAGE and visualized by fluorescence imaging.

### Analysis of cGAS reaction order

To investigate the catalytic mechanism of *S. marcescens* cGAS, reactions containing 2 µM cGAS were supplemented with 250 µM non-hydrolysable nucleotide analogues Ap(c)pp and Gp(c)pp (Jena Bioscience) in place of ATP and GTP. Reactions were incubated at 37 °C for 30 min, heat-inactivated at 80 °C for 10 min and analysed by HPLC.

To analyse the second catalytic step, reactions containing 250 µM 5′-pppA[2′–5′]pG, 5′-pppA[3′–5′]pG or 5′-pppG[2′–5′]pA were supplemented with 2 µM cGAS. Following incubation at 37 °C for 15 min, aliquots were collected for analysis before and after nuclease P1 digestion. The remaining reaction mixture was supplemented with 2 µM Exo and incubated at 37 °C for an additional 1 h before heat inactivation and HPLC analysis.

### HPLC analysis

Reaction mixtures were clarified by centrifugation at 21,000 × g for 15 min before analysis. Standard HPLC analyses were performed using an Agilent 1260 Infinity II LC system equipped with a Luna Omega 5 µm Polar C18 100 Å column (150 × 4.6 mm; Phenomenex). Samples (10 µL) were injected at a flow rate of 1.0 mL min^−1^ using solvent A (acetonitrile; Sigma-Aldrich) and solvent B (20 mM ammonium acetate; Sigma-Aldrich). The elution program was: 0–5 min, 100% B; 5–20 min, 100–70% B; 20–23 min, 60% B; followed by re-equilibration at 100% B. Elution profiles were monitored by UV absorbance at 254 nm.

For detection of reaction intermediates, analyses were performed at a flow rate of 0.5 mL min^−1^ using the following gradient: 0–10 min, 100% B; 10–25 min, 100–85% B; 25–26 min, 85–60% B; 26–31 min, 60% B; 31–33 min, 60–100% B; followed by 7 min re-equilibration at 100% B.

Peak areas were converted to molar quantities using calibration curves generated with chemically synthesized standards, including 3′,2′-cGAMP (Biolog), 2′,3′-cGAMP (Biolog), 5′-pppA[2′–5′]pG, AMP, GMP and GTP. Data were analysed using GraphPad Prism v8.0.2.

### Competition assays

To investigate competition between cGAS and Exo for exogenously supplied intermediates, reactions containing 25 µM 5′-pppA[2′–5′]pG were supplemented with 2 µM cGAS and either 20 nM or 2 µM Exo. Additional reactions containing 250 µM 5′-pppA[2′–5′]pG were performed under identical conditions. Reactions were incubated at 37 °C for 15 min, heat-inactivated at 80 °C for 10 min and analysed by HPLC.

To assess competition for intermediates generated de novo by cGAS, reactions containing 25 µM ATP and GTP were supplemented with 2 µM cGAS alone or together with Exo at the indicated concentrations. Reactions lacking enzyme served as negative controls. Products were analysed by HPLC following incubation at 37 °C for 15 min and heat-inactivated at 80 °C for 10 min.

### Multiple sequence alignments

Multiple sequence alignments were generated using Clustal Omega and visualized using Jalview v2.11.2.5. Alignments included *S. marcescens* Exo, *E. coli* upec-117 Cap18, *V. cholerae* Exo, the ε subunit of *E. coli* DNA polymerase III (PDB: 5FKU_D), *Pseudomonas aeruginosa* RNase T (MBG7179145.1) and *V. cholerae* oligoribonuclease (WP_000010183.1). Alignments were performed using the BLOSUM substitution matrix with gap opening and extension penalties of 10 and 0.05, respectively.

### Phylogenetic analysis

Exo homolog sequences were obtained by searching UniRef30 database using HH-suite (doi: 10.1186/s12859-019-3019-7). Polymerase subunits were manually removed from the dataset. Sequences were clustered using MMseqs2 with a threshold of 0.3 identity, generating 492 sequences. The structures of Exo homologs were predicted using Alphafold3. We then computed AA and 3Di MSTAs using FoldMason. MSTAs were trimmed using trimAl, removing columns containing ≥35% gaps. We used iqtree2 to reconstruct a phylogenic tree with 1000 bootstrap replicate and substitution model PMB+F+R10. Tree visualization was performed using iTol.

### Protein crystallization and structure determination

The *V. cholerae* Exo gene was codon-optimized and cloned into pET28a-HT encoding an N-terminal hexahistidine tag followed by a TEV protease cleavage site. Protein expression was performed in *E. coli* BL21(DE3) grown in LB medium supplemented with 50 µg mL^−1^ kanamycin. Cultures were induced at an OD600 of 0.6 with 0.25 mM IPTG and incubated overnight at 16 °C.

Cells were harvested by centrifugation and resuspended in lysis buffer containing 20 mM Tris-HCl, pH 8.0, 250 mM NaCl, 1 mM TCEP, 20 mM imidazole and 5% glycerol supplemented with 1 mM PMSF. Cells were lysed by sonication and clarified by centrifugation at 21,000 × g for 1 h at 4 °C. Clarified lysates were filtered through 0.22 µm filters and incubated with Ni–NTA resin for 1 h at 4 °C. Following washing and elution, TEV protease was added and samples were incubated overnight at 4 °C. Cleaved proteins were subjected to reverse Ni–NTA purification, followed by anion-exchange chromatography on a HiTrap Capto Q column (GE Healthcare) using a linear gradient of 50–500 mM NaCl in 20 mM Tris-HCl, pH 8.8, and 10 mM β-mercaptoethanol. Proteins were further purified by size-exclusion chromatography on a Superdex 200 Increase 10/300 column (GE Healthcare) equilibrated in 20 mM Tris-HCl, pH 8.0, 150 mM NaCl and 1 mM TCEP. Peak fractions were concentrated to 10 mg mL^−1^ and stored in 20 mM Tris-HCl, pH 8.0, 50 mM NaCl and 1 mM TCEP at −80 °C.

Phosphorothioate A[2′–5′]psG and A[2′–5′]pG ligands were synthesized by GenScript. Exo–ligand complexes were formed by incubation at a 1:1.5 molar ratio for 30 min at 4 °C. Crystallization screening was performed at 20 °C using sitting-drop vapour diffusion with commercial sparse-matrix screens. Optimized crystals were obtained by hanging-drop vapour diffusion in 0.25 M NaSCN and 24% PEG 3350.

Crystals were cryoprotected with reservoir solution supplemented with 20% glycerol and flash-cooled in liquid nitrogen. Diffraction data were collected at beamline BL10U2 of the Shanghai Synchrotron Radiation Facility (SSRF) to resolutions of 1.96 Å for the phosphorothioate A[2′–5′]psG complex and 1.99 Å for the hydrolysable complex. Structures were solved by molecular replacement in PHENIX using AlphaFold2-predicted Exo models as search templates, manually rebuilt in Coot and refined iteratively in PHENIX. Coordinates and structure factors were deposited in the Protein Data Bank under accession codes 22HD and 22HE. Data collection and refinement statistics are summarized in Extended Data Table 1.

Structural figures were generated using UCSF ChimeraX v1.10.1.

### Structure prediction

Structures of *S. marcescens* Exo were predicted using AlphaFold3. The top-ranked model was used for visualization and comparisons in PyMOL.

### Plaque assays

Phages were amplified in *E. coli* B cultures at multiplicities of infection ranging from 1:100 to 1:1,000 and incubated at 37 °C until complete lysis. Lysates were clarified by centrifugation and filtered through 0.22 µm filters.

Small-drop plaque assays were performed as previously described. Wild-type or mutant E2-CBASS operons were cloned into pQE82L and transformed into *E. coli* BL21. Empty pQE82L served as control. Cultures were grown to OD600 ∼0.2, induced with 50 µM IPTG and incubated for 1 h until OD600 reached 0.6–0.7. Cells (800 µL) were mixed with 25 mL molten LB top agar containing 0.75% agar, antibiotics and IPTG before pouring onto LB agar plates. Serially diluted phage stocks were spotted onto plates, which were incubated at 37 °C for 16–18 h before imaging.

### mRNA synthesis and purification

Modified linear WT or mutant Exo mRNAs encoded bacterial Exo proteins and contained a FLAG tag, human β-globin 5′ and 3′ untranslated regions, a Cap 1 structure and a 120-nucleotide poly(A) tail. mRNAs were synthesized in vitro from linearized DNA templates using T7 RNA polymerase-mediated (New England Biolabs) transcription with complete substitution of uridine triphosphate by pseudouridine triphosphate (ΨTP; Apexbio). Following transcription, reactions were treated with DNase I (Promega) for 15 min and purified using the Monarch RNA Cleanup Kit (New England Biolabs) according to the manufacturer’s instructions. RNA integrity was verified by agarose gel electrophoresis before use.

### Cell culture

HEK293T and RAW264.7 cells were maintained in DMEM supplemented with 10% fetal bovine serum, 100 U mL^−1^ penicillin and 100 µg mL^−1^ streptomycin at 37 °C in a humidified atmosphere containing 5% CO2. Cell lines were routinely tested and confirmed negative for mycoplasma contamination.

### Cytotoxicity assay

RAW264.7 cells were treated with Exo mRNA at the indicated concentrations, while control cells received culture medium alone. After 12 h, medium was replaced with 100 µL fresh medium containing 10 µL Cell Counting Kit-8 reagent (Dojindo), followed by incubation for 1.5 h at 37 °C. Absorbance at 450 nm was measured using a Cytation 5 microplate reader (BioTek). Cell viability was normalized to untreated controls.

### cGAMP ELISA

RAW264.7 cells (1 × 10^6^ cells per well) were seeded in six-well plates and transfected with 1 µg Exo mRNA. After 12 h, cells were stimulated by transfection with HT-DNA at a final concentration of 1 µg mL^−1^ for 1 h. Intracellular cGAMP levels were quantified using a commercial ELISA kit (Cayman) according to the manufacturer’s instructions.

### RNA extraction and Real-Time PCR (RT-PCR)

Total RNA was extracted from cultured cells or mouse tissues and reverse-transcribed into cDNA. Quantitative PCR was performed using SYBR Green Master Mix on a QuantStudio 3 real-time PCR system (Thermo Fisher Scientific). Relative transcript levels were calculated using the ΔΔCt method and normalized to *Gapdh* expression.

Primer sequences were as follows:

*Gapdh* forward, 5′-CATCACTGCCACCCAGAAGACTG-3′; reverse, 5′-ATGCCAGTGAGCTTCCCGTTCAG-3′

*Ifnb* forward, 5′-TCCTGCTGTGCTTCTCCACCACA-3′; reverse, 5′-AAGTCCGCCCTGTAGGTGAGGTT-3′

*Exo* forward, 5′-GATCTGGACGAGCTGACAAGGAGC-3′; reverse, 5′-AACCGAAGGGATTCACGCCAAAGAA-3′

*Il-6* forward, 5′-CTGCAAGAGACTTCCATCCAG-3′; reverse, 5′-AGTGGTATAGACAGGTCTGTTGG-3′

*Il1b* forward, 5′-TCTCAGATTCACAACTGTTCGTG-3′; reverse, 5′-AGAAAATGAGGTCGGTCTCACTA-3′

### Immunoblotting

Cells were lysed in RIPA buffer (Beyotime) supplemented with EDTA-free protease inhibitor cocktail. Protein concentrations were determined using BCA assays (Beyotime). Equal amounts of protein were resolved by SDS–PAGE and transferred onto nitrocellulose membranes. Membranes were blocked with 5% non-fat milk for 1 h at room temperature and incubated overnight at 4 °C with primary antibodies, followed by incubation with horseradish peroxidase-conjugated secondary antibodies for 1 h at room temperature. Signals were detected using a MiniChemi 610 imaging system (Sinsage). Primary antibodies used were: STING (Proteintech), phospho-STING (Cell Signaling Technology) and GAPDH (Abcam).

### Animal studies

All animal experiments were approved by the Animal Care and Use Committee of the Guangzhou National Laboratory and conducted in accordance with institutional guidelines for animal care and use. Mice were housed under specific pathogen-free conditions at 23 ± 2 °C with 50 ± 20% humidity under a 12 h light–dark cycle.

### Androgenetic alopecia mouse model

Six-week-old male C57BL/6 mice were acclimated for one week before random assignment into experimental groups. To induce androgenetic alopecia, mice received subcutaneous injections of testosterone propionate (3 mg per mouse) every other day for 14 d. Control mice received corn oil vehicle injections on the same schedule.

Following testosterone treatment, dorsal hair was removed by shaving followed by depilatory cream application. Mice subsequently received weekly subcutaneous injections of lipid nanoparticle-formulated Exo mRNA (10 µg) or empty lipid nanoparticles as vehicle controls, while testosterone administration continued throughout the treatment period. Hair regrowth was documented photographically at the indicated time points and quantified using ImageJ software. Histological analyses were performed on dorsal skin sections collected at the experimental endpoint.

Investigators were blinded during quantitative analysis of hair regrowth and histological assessment.

### Statistical analysis

All experiments were independently repeated at least three times unless otherwise stated. Data are presented as mean ± s.d. Statistical analyses were performed using GraphPad Prism 10. Two-tailed unpaired Student’s *t*-tests were used for comparisons between two groups. One-way ANOVA followed by Tukey’s multiple-comparison test was used for comparisons involving more than two groups unless otherwise indicated. Exact sample sizes and statistical details are provided in the figure legends. Differences were considered statistically significant at *P* < 0.05.

---

**Fig. S1.**
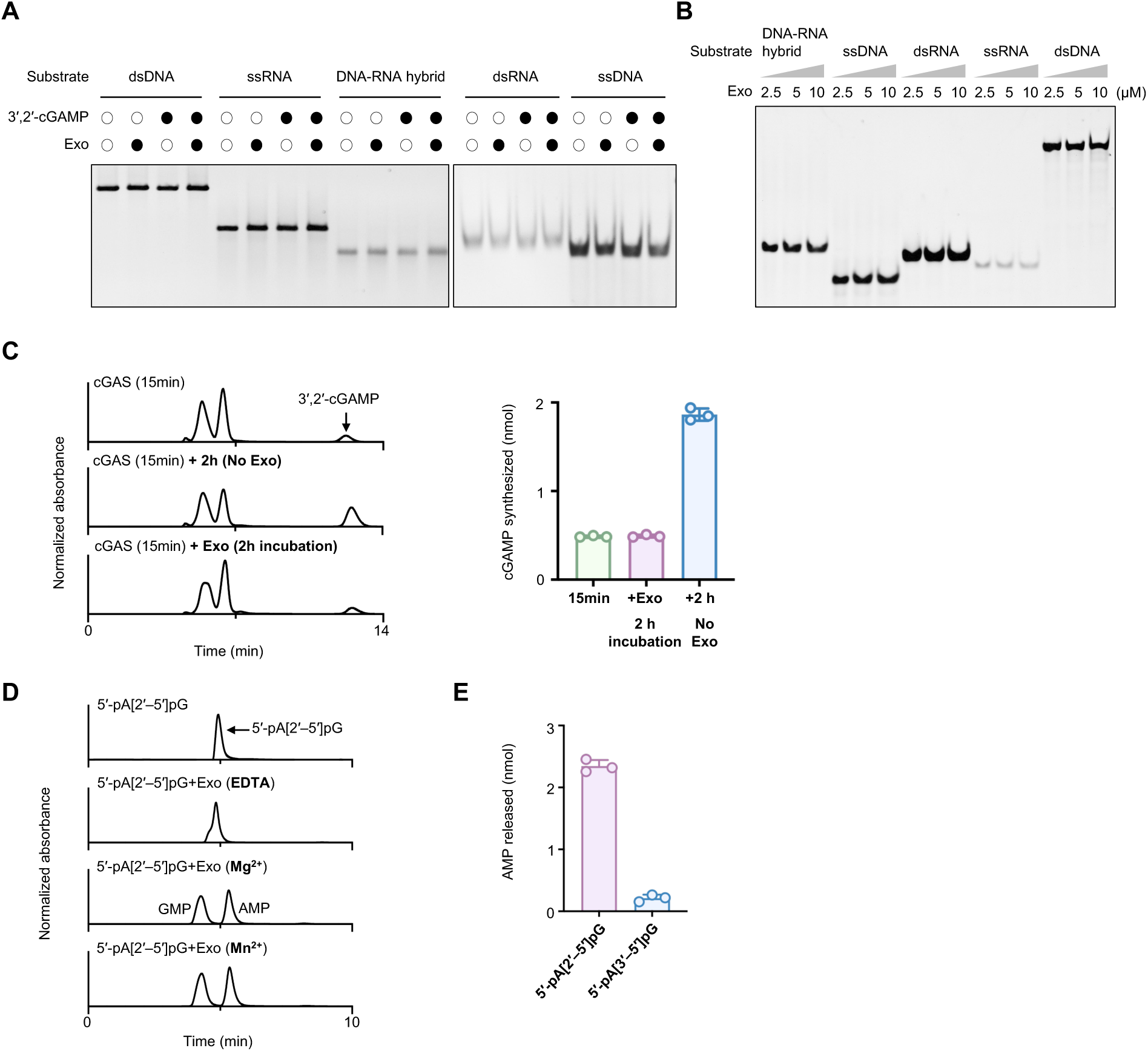
Exo lacks canonical nuclease activity but targets catalytic intermediates of cGAS. (**A**) Agarose gel electrophoresis analysis of nucleic acid degradation assays. *Sm*Exo does not degrade dsDNA, ssDNA, dsRNA, ssRNA or DNA–RNA hybrids under conditions permissive for exonuclease activity. (**B**) Electrophoretic mobility shift assays (EMSAs) using increasing concentrations of *Sm*Exo and nucleic acid substrates. No mobility shift is observed. (**C**) HPLC analysis and quantification of 3′,2′-cGAMP accumulation during ongoing *S. marcescens* cGAS reactions. Addition of *Sm*Exo prevents further accumulation of cGAMP without reducing preformed product. Data are mean ± s.d., n = 3 independent experiments. (**D**) Divalent metal dependence of Exo activity. Hydrolysis of 5′-pA[2′–5′]pG occurs in the presence of Mg²⁺ or Mn²⁺ but not in the absence of divalent cations. (**E**) Quantification of Exo activity toward 5′-pA[2′–5′]pG and 5′-pA[3′–5′]pG. Data are mean ± s.d., n = 3 independent experiments.

**Fig. S2.**
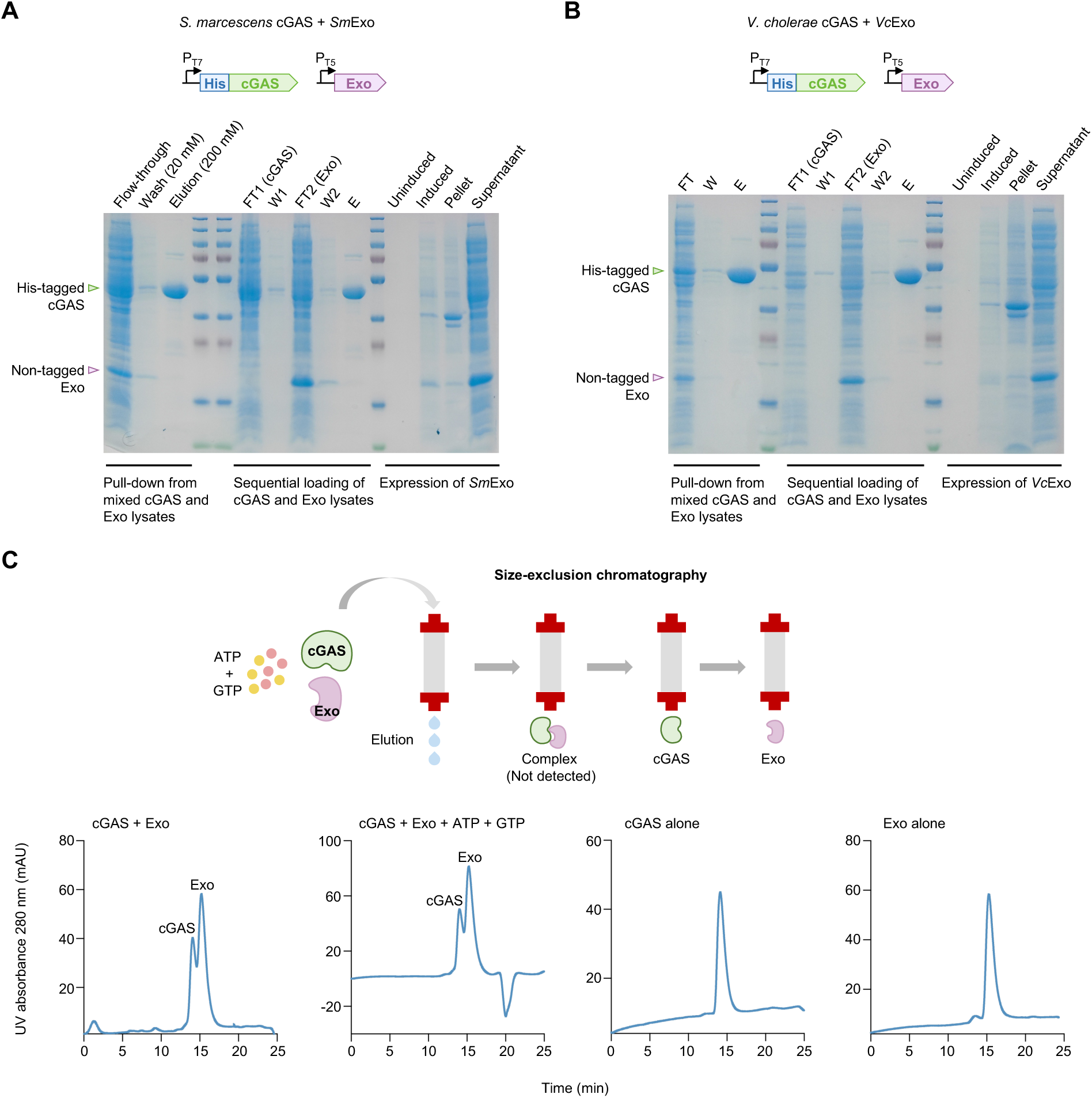
Exo does not regulate cGAS through direct interaction. (**A**) Pull-down assays testing interaction between *Sm*Exo and *S. marcescens* cGAS. (**B**) Pull-down assays testing interaction between *Vc*Exo and *V. cholerae* cGAS. (**C**) Size-exclusion chromatography analysis of *Vc*Exo and *V. cholerae* cGAS in the presence or absence of ATP and GTP subtrates.

**Fig. S3.**
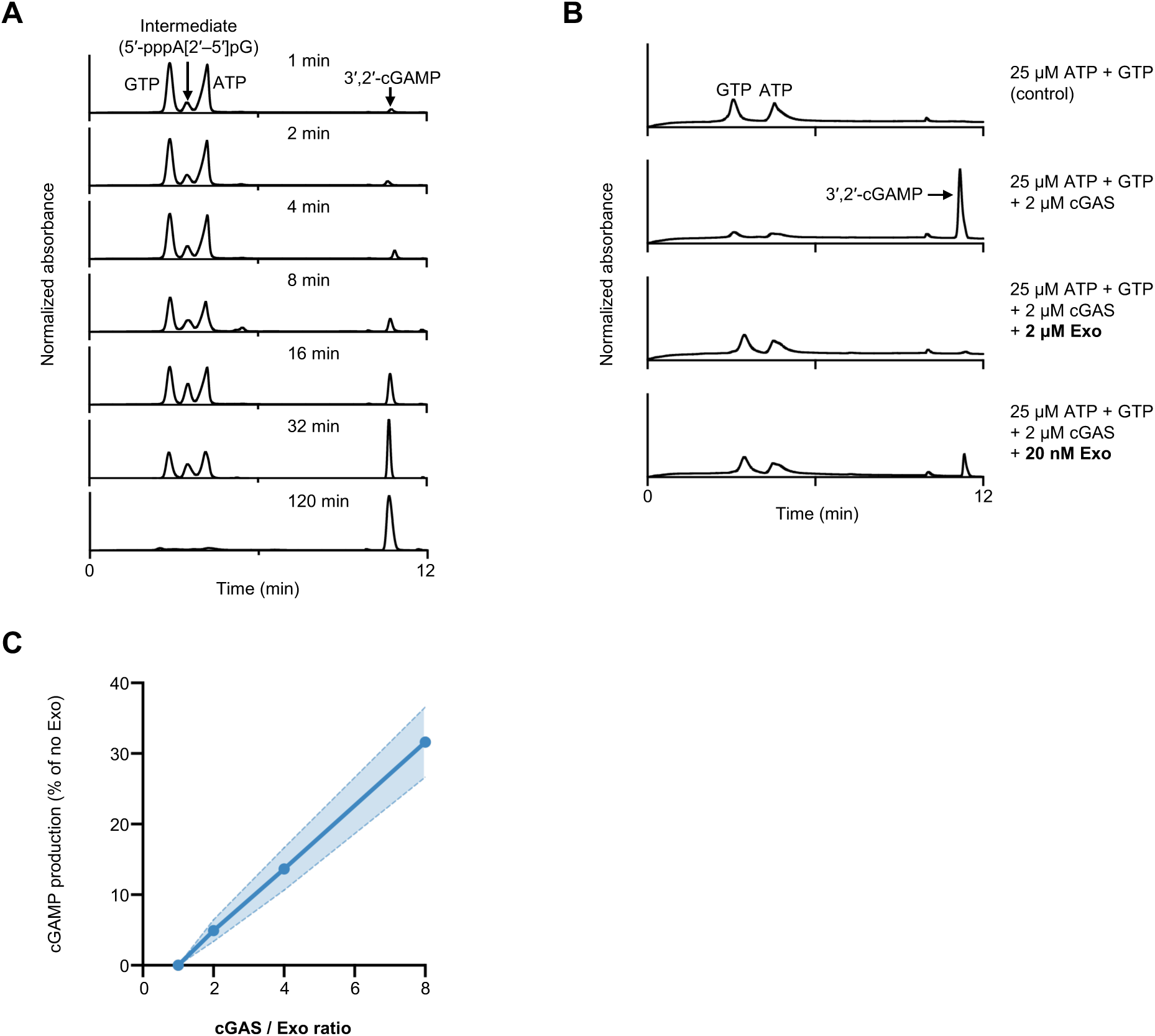
Catalytic intermediate turnover defines a threshold checkpoint. (**A**) HPLC analysis of 3′,2′-cGAMP synthesis by *S. marcescens* cGAS at the indicated time points. (**B**) HPLC analysis showing depletion of substrates and accumulation of cyclization products in the reactions containing cGAS, Exo and the ATP and GTP substrates under the indicated concentrations. (**C**) Quantification of 3′,2′-cGAMP production at varying cGAS:Exo ratios. Data are mean ± s.d., n = 3 independent experiments.

**Fig. S4.**
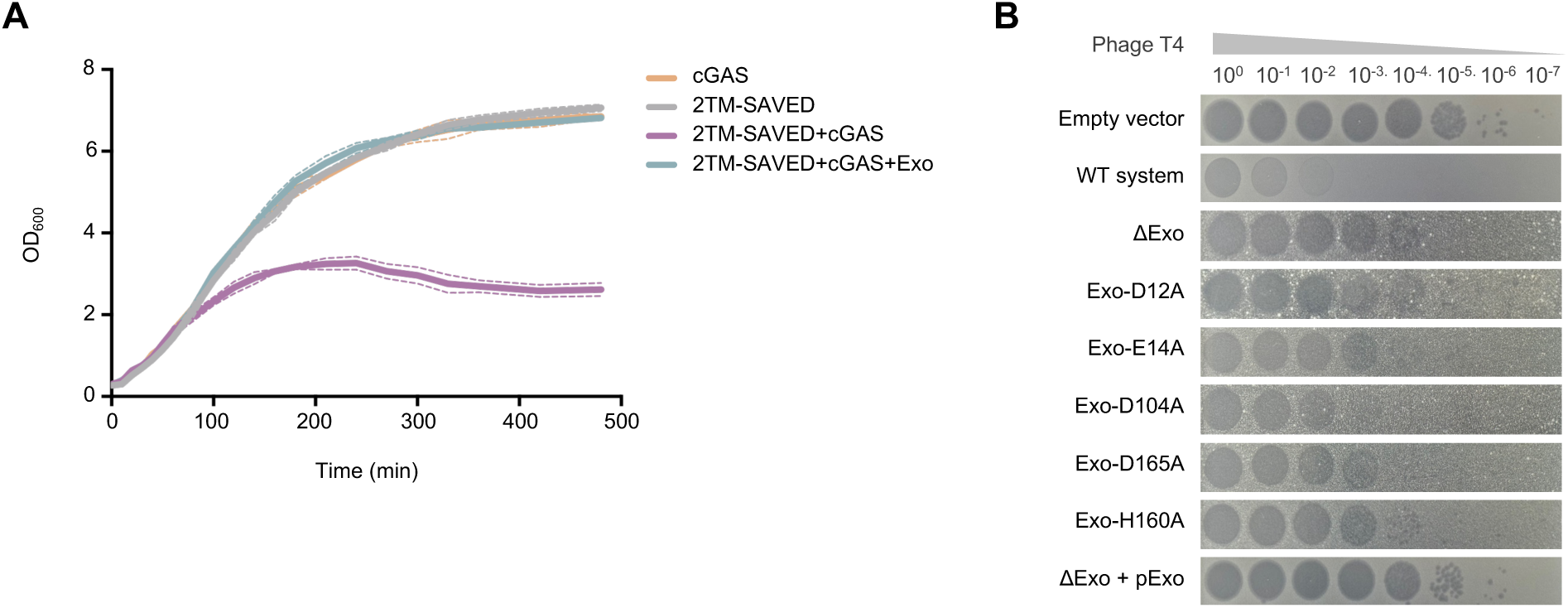
Exo restrains CBASS activation in vivo. (**A**) Bacterial growth curves showing that expression of cGAS or 2TM-SAVED alone does not impair growth, whereas co-expression results in cytotoxicity. Co-expression of Exo suppresses this phenotype. (**B**) Phage T4 infection assays in cells expressing an empty vector, the wild-type E2-CBASS operon, an Exo-deficient operon, or operons carrying Exo mutations. Data are representative of three independent experiments.

**Fig. S5.**
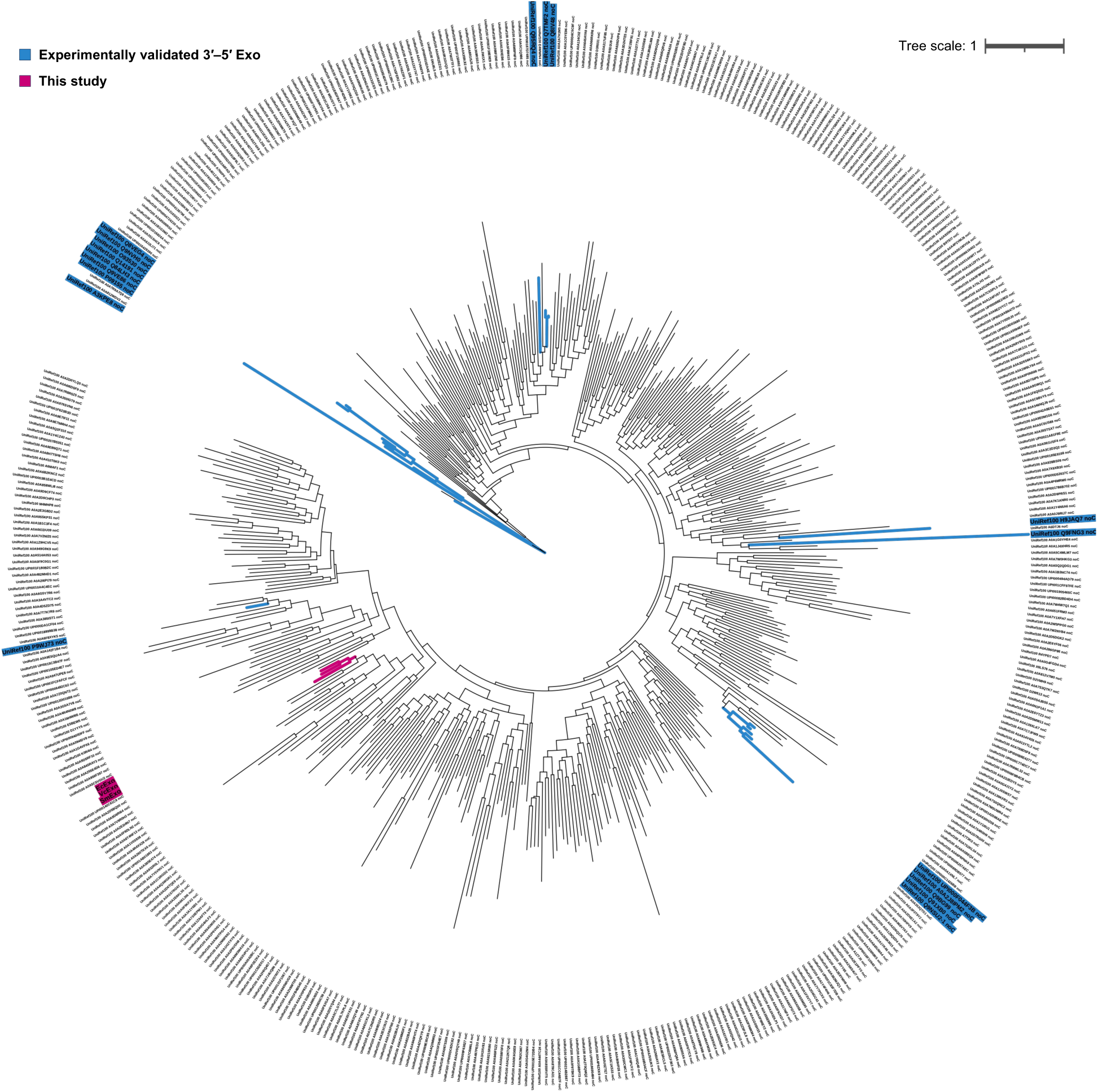
Exo forms a distinct clade within DEDDh exonucleases. Phylogenetic tree of DEDDh family exonucleases showing that CBASS-associated Exo proteins form a distinct clade. Exo homologues analysed in this study are highlighted, and experimentally validated 3′–5′ exonucleases are indicated for comparison.

**Fig. S6.**
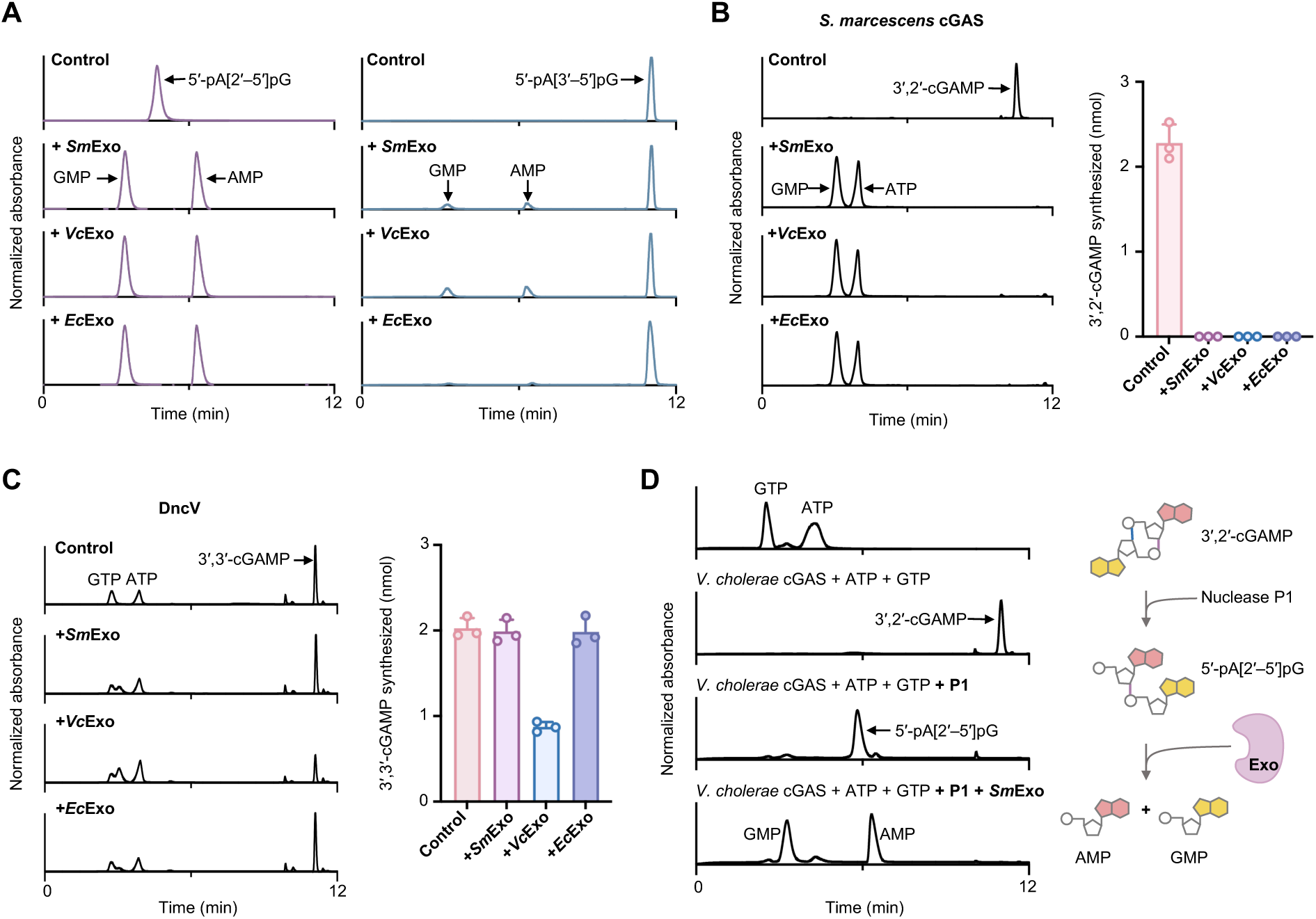
Exo selectively targets 2′–5′-linked cyclic dinucleotide intermediates. (**A**) HPLC analysis of hydrolysis of 5′-pG[2′–5′]pA and 5′-pG[3′–5′]pA by Exo homologues from *S. marcescens*, *V. cholerae* and *E. coli* upec-117. (**B**) HPLC analysis and quantification of 3′,2′-cGAMP synthesis by *S. marcescens* cGAS in the presence or absence of Exo homologues. Data are mean ± s.d., n = 3 independent experiments. (**C**) HPLC analysis and quantification of 3′,3′-cGAMP synthesis by *V. cholerae* DncV in the presence or absence of Exo homologues. Data are mean ± s.d., n = 3 independent experiments. (**D**) HPLC analysis of cyclic dinucleotide products synthesized by the *V. cholerae* CBASS-associated cGAS under the indicated conditions.

**Fig. S7.**
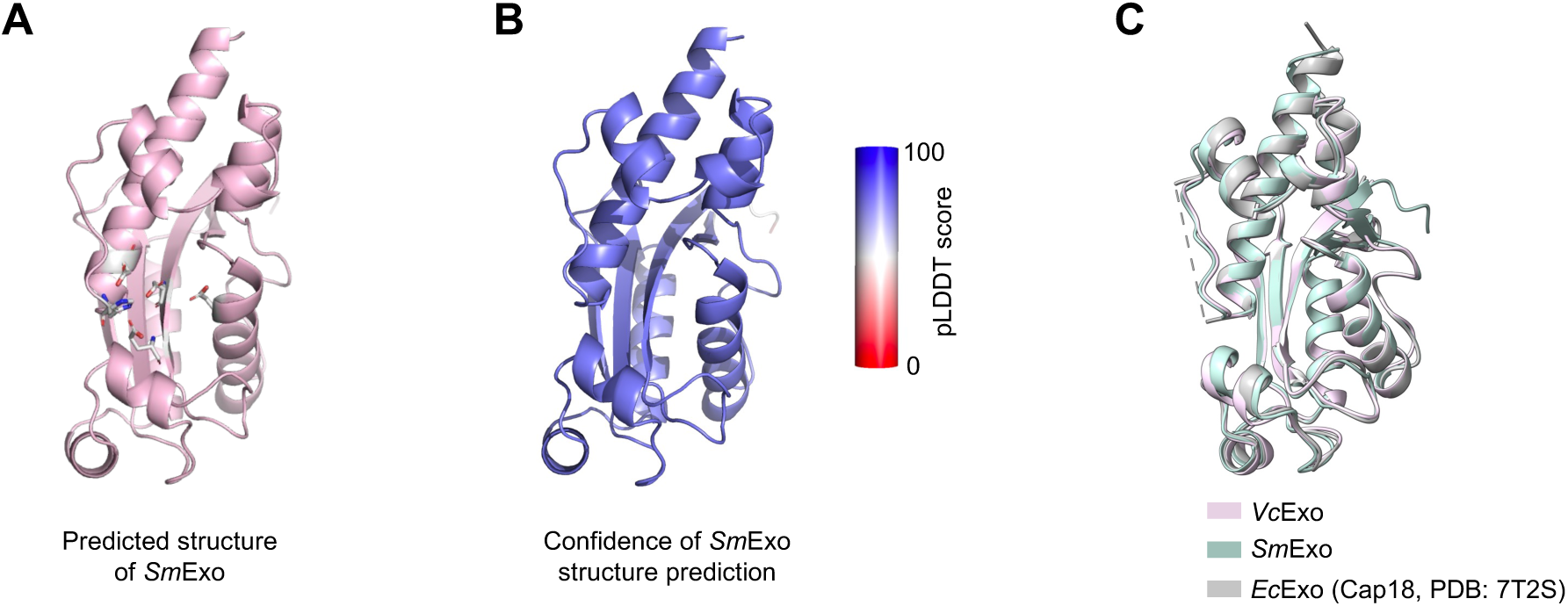
Structural conservation of Exo proteins. (**A**) AlphaFold-predicted structure of *Sm*Exo with catalytic residues indicated. (**B**) *Sm*Exo structure coloured by per-residue confidence (pLDDT). (**C**) Structural comparison of *Vc*Exo (pink) with the predicted structure of *Sm*Exo (green) and the crystal structure of *Escherichia coli* upec-117 Exo (*Ec*Exo, grey), revealing a highly conserved overall fold among CBASS-associated Exo homologues.

**Fig. S8.**
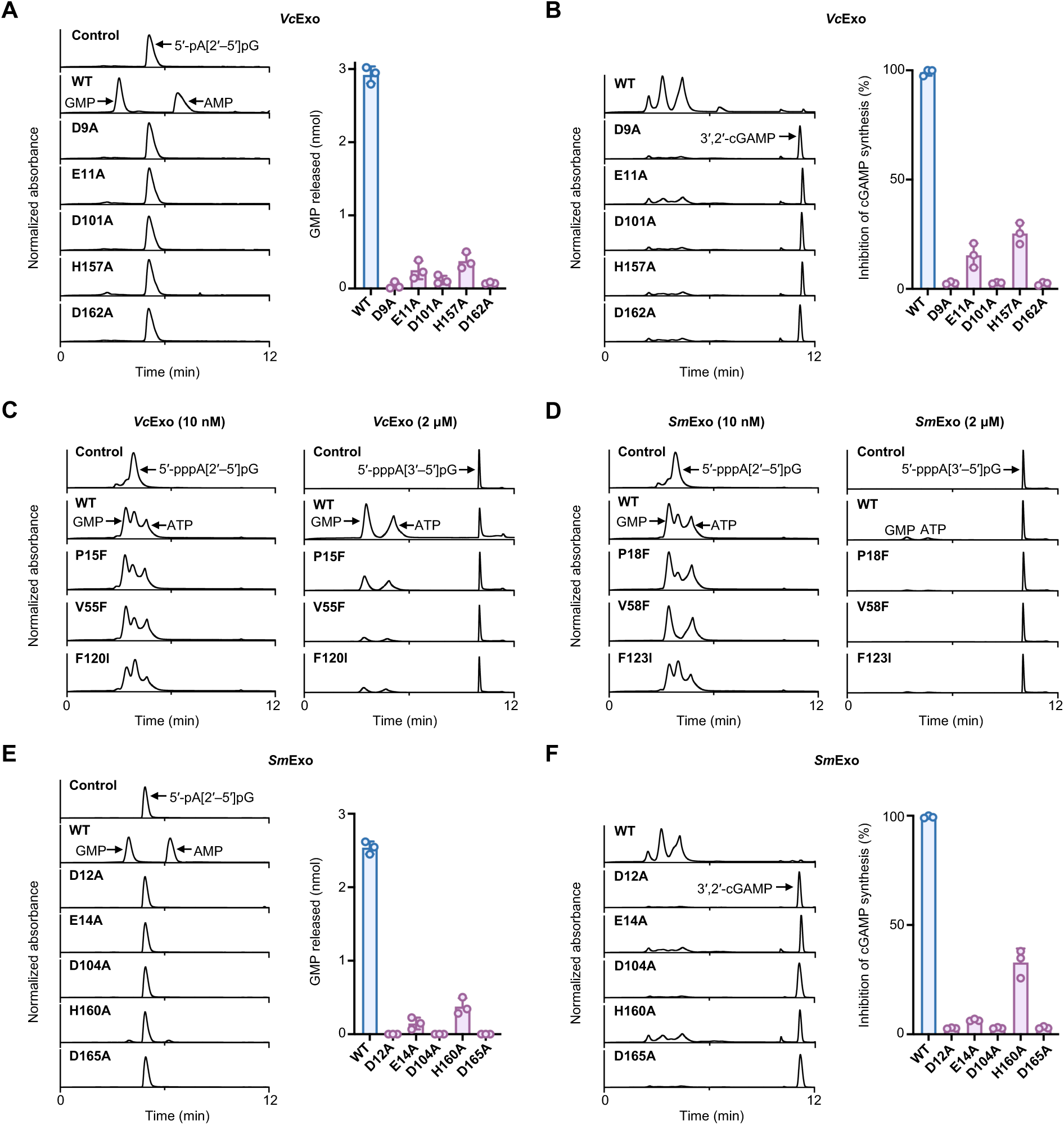
Structural determinants of 2′–5′ linkage selectivity. (**A**) HPLC analysis of degradation of the 5′-pA[2′–5′]pG substrate by *Vc*Exo catalytic mutants. Data are mean ± s.d., n = 3 independent experiments. (**B**) HPLC analysis of *V. cholerae* cGAS activity in the presence of *Vc*Exo mutants. Data are mean ± s.d., n = 3 independent experiments. (**C**) HPLC analysis of cleavage of 5′-pppA[2′–5′]pG and 5′-pppA[3′–5′]pG by *Vc*Exo variants. (**D**) HPLC analysis of cleavage of 5′-pppA[2′–5′]pG and 5′-pppA[3′–5′]pG by *Sm*Exo variants. (**E**) HPLC analysis of degradation of 5′-pA[2′–5′]pG by *Sm*Exo catalytic mutants. Data are mean ± s.d., n = 3 independent experiments. (**F**) HPLC analysis of *S. marcescens* cGAS activity in the presence of *Sm*Exo mutants. Data are mean ± s.d., n = 3 independent experiments.

**Fig. S9.**
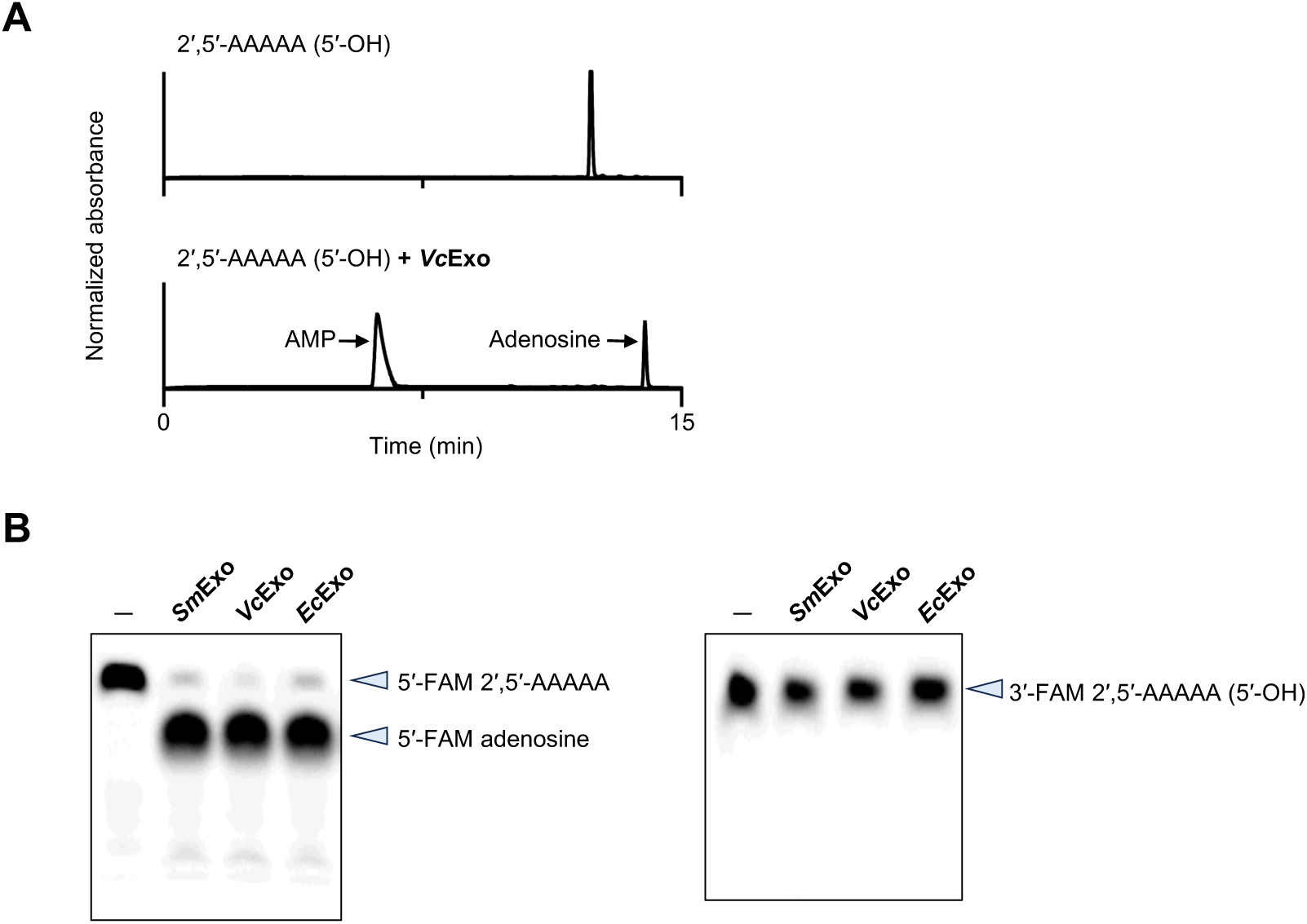
Exo cleaves 2′–5′-linked oligoadenylate molecules. (**A**) HPLC analysis of Exo-mediated cleavage of a 2′–5′-linked oligoadenylate (2′–5′-AAAAA). (**B**) Polyacrylamide gel electrophoresis analysis of cleavage of 2′–5′-AAAAA substrates labelled at the 5′ or 3′ terminus.

**Fig. S10.**
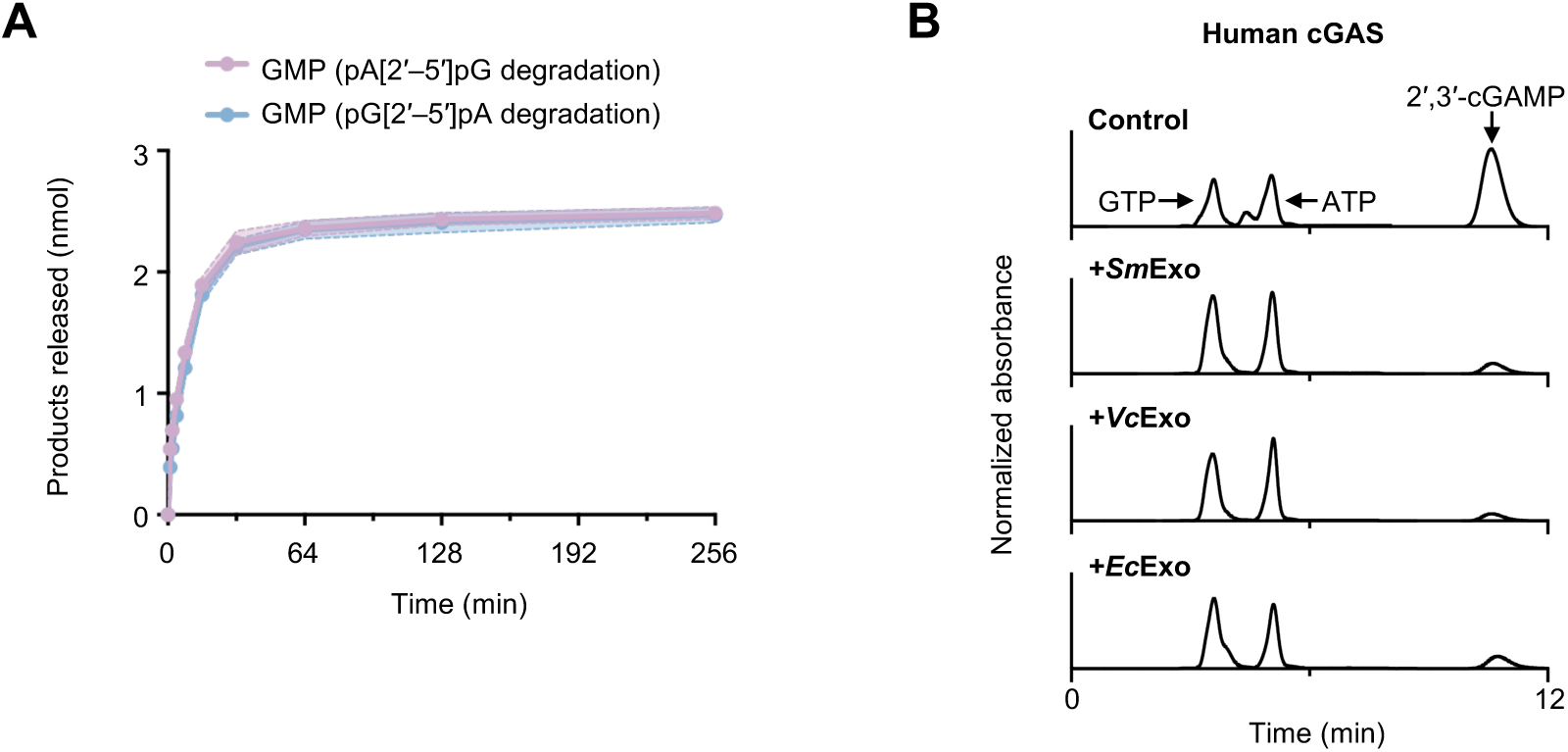
Exo targets catalytic intermediates of human cGAS. (**A**) Quantification of Exo-mediated hydrolysis of 5′-pA[2′–5′]pG and 5′-pG[2′–5′]pA substrates, showing comparable cleavage efficiencies for both dinucleotides. (**B**) HPLC analysis of 2′,3′-cGAMP synthesis by human cGAS in the presence or absence of Exo homologues.

**Fig. S11.**
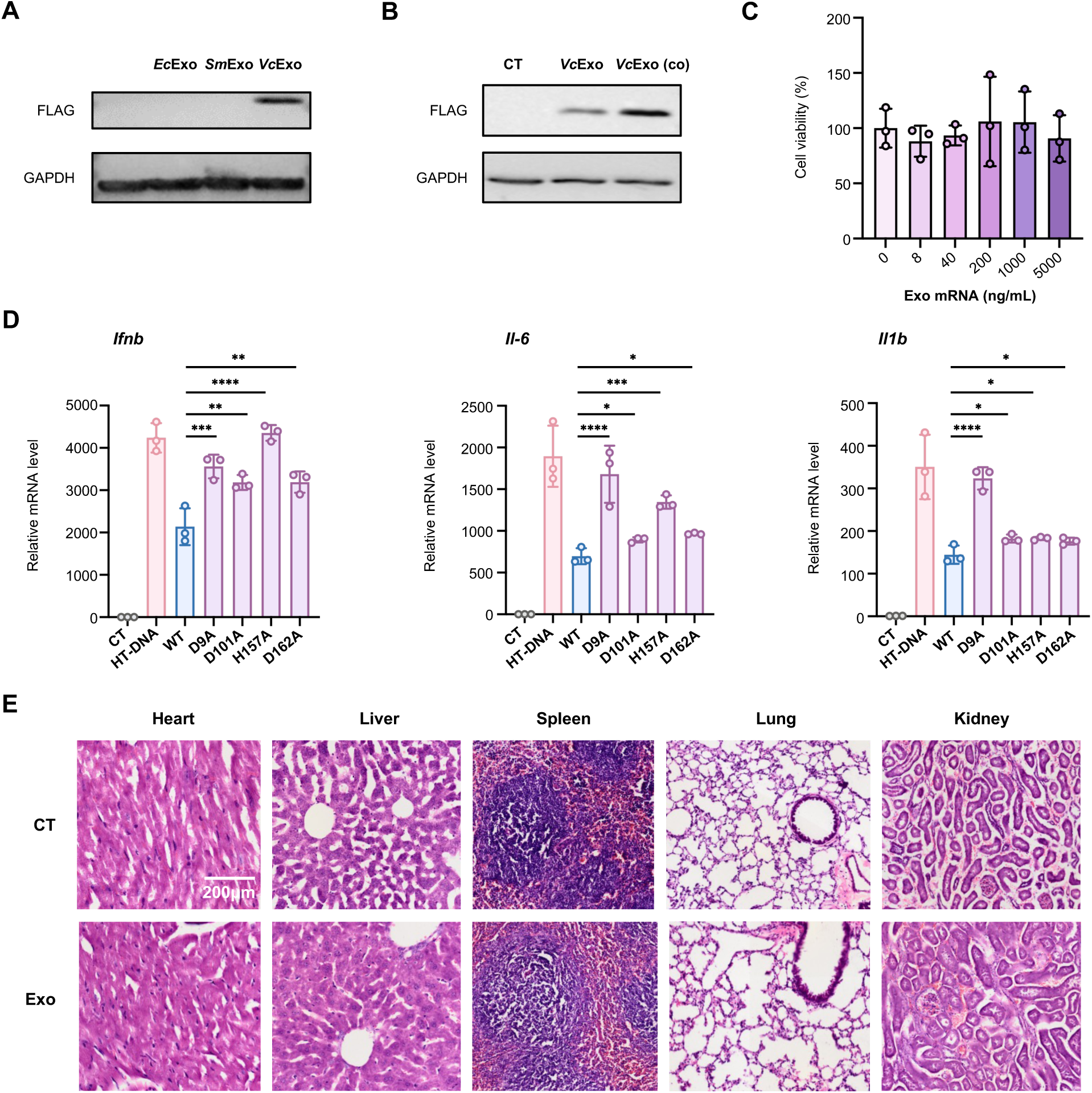
Exo suppresses cGAS signalling in mammalian systems. (**A**) Immunoblot analysis of FLAG-tagged Exo proteins expressed in HEK293T cells. (**B**) Immunoblot analysis of *Vc*Exo expressed from native or human codon-optimized (co) mRNA. (**C**) Cell viability of RAW264.7 cells treated with Exo mRNA. Data are mean ± s.d., n = 3 independent experiments. (**D**) RT–PCR analysis of inflammatory gene interferon-β (*Ifnb*), interleukin-6 (*Il-6*) and interleukin-1β (*Il1b*) expression in HT-DNA–stimulated RAW264.7 cells transfected with Exo variants. Data are mean ± s.d., n = 3 independent experiments. (**E**) Representative H&E stained sections of major organs from treated mice.

**Table S1.** Data collection and refinement statistics for the *V. cholerae* Exo with phosphorothioate A[2′–5′]psG or A[2′–5′]pG crystals.

|  | *V. cholerae* Exo with phosphorothioate<br>A[2'–5']psG | *V. cholerae* Exo with A[2'–<br>5']pG |
| --- | --- | --- |
| **Data collection** |  |  |
| Space group | C 1 2 1 | C 1 2 1 |
| Cell dimensions |  |  |
| *a*, *b*, *c* (Å) | 95.36, 59.48, 187.83 | 95.25, 59.57, 186.80 |
| $\alpha$, $\beta$, $\gamma$ (°) | 90.0, 91.899, 90.0 | 90.0, 91.95, 90.0 |
| Resolution (Å) | 62.58 – 1.96 | 25.25 – 1.99 |
| *R*<sub>merge</sub> | 0.051 (0.128) | 0.112 (0.294) |
| *I* / $\sigma I$ | 23.800 (9.900) | 12.200 (5.100) |
| Completeness (%) | 98.9 (90.9) | 98.7 (97.4) |
| Redundancy | 5.8 (4.0) | 5.8 (4.5) |
| **Refinement** |  |  |
| Resolution (Å) | 62.58 – 1.96 | 93.36 – 2.00 |
| No. reflections | 74862 | 71095 |
| *R*<sub>work</sub> / *R*<sub>free</sub> | 0.1607 / 0.2095 | 0.1793 / 0.2418 |
| No. atoms |  |  |
| Protein | 8129 | 8268 |
| Ligand | 138 | 98 |
| Water | 1039 | 886 |
| Average *B*-factor | 21.26 | 24.70 |
| R.m.s. deviations |  |  |
| Bond lengths (Å) | 0.006 | 0.007 |
| Bond angles (°) | 0.79 | 0.81 |

